# Transcriptional alterations in opioid use disorder reveal an interplay between neuroinflammation and synaptic remodeling

**DOI:** 10.1101/2020.09.14.296707

**Authors:** Marianne L. Seney, Sam-Moon Kim, Jill R. Glausier, Mariah A. Hildebrand, Xiangning Xue, Wei Zong, Jiebiao Wang, Micah A. Shelton, BaDoi N. Phan, Chaitanya Srinivasan, Andreas R. Pfenning, George C. Tseng, David A. Lewis, Zachary Freyberg, Ryan W. Logan

## Abstract

**Background:** Prevalence rates of opioid use disorder (OUD) have increased dramatically, accompanied by a surge of overdose deaths. While opioid dependence has been extensively studied in preclinical models, an understanding of the biological alterations that occur in the brains of people who chronically use opioids and who are diagnosed with OUD remains limited. To address this limitation, RNA-sequencing (RNA-seq) was conducted on the dorsolateral prefrontal cortex (DLPFC) and nucleus accumbens (NAc), regions heavily implicated in OUD, from postmortem brains in subjects with OUD.

**Methods:** We performed RNA-seq on the DLPFC and NAc from unaffected comparison subjects (n=20) and subjects diagnosed with OUD (n=20). Our transcriptomic analyses identified differentially expressed (DE) transcripts and investigated the transcriptional coherence between brain regions using rank-rank hypergeometric ordering (RRHO). Weighted gene co-expression analyses (WGCNA) also identified OUD-specific modules and gene networks. Integrative analyses between DE transcripts and GWAS datasets using linkage disequilibrium score (LDSC) assessed the genetic liability psychiatric-related phenotypes.

**Results:** RRHO analyses revealed extensive overlap in transcripts between DLPFC and NAc in OUD, primarily relating to synaptic remodeling and neuroinflammation. Identified transcripts were enriched for factors that control pro-inflammatory cytokine-mediated, chondroitin sulfate, and extracellular matrix signaling. Cell-type deconvolution implicated a role for microglia as a critical driver for opioid-induced neuroplasticity. Using LDSC, we discovered genetic liabilities for risky behavior, attention deficit hyperactivity disorder, and depression.

**Conclusions:** Overall, our findings reveal new connections between the brain’s immune system and opioid dependence in the human brain.

## Introduction

Prevalence of opioid use disorder (OUD) and deaths from opioid overdose have soared in the United States (1). The enormity of the public health impact have been the impetus for broad efforts to develop new treatments for OUD. Progress towards effective therapeutics requires better understanding of the alterations in the brain of those who develop dependence.

Impulsivity and deficits in cognition are hallmarks of OUD (3). These impairments have been attributed to functional alterations in corticostriatal circuits including the dorsolateral prefrontal cortex (DLPFC) and nucleus accumbens (NAc) (3, 4). Moreover, a history of substance use is associated with corticostriatal circuit dysfunction that contributes to cognitive impairment and promotes risky behavior (5). However, we still have a limited understanding of the cellular and molecular alterations due to chronic opioid use and OUD that occur in these circuits in human brain.

Although few studies have studied postmortem brains in subjects diagnosed with OUD, the approach has potential to uncover relevant and therapeutically viable pathways in the brain in opioid dependence. Previous work reported changes in opioid receptor expression in DLPFC (6–8) and altered expression of the machinery that regulates presynaptic glutamate release in NAc (9, 10), potentially related to addiction severity in heroin users. Preclinical evidence has corroborated these findings by demonstrating unique interactions between opioid and glutamate receptor signaling in opioid withdrawal and dependence (11–13). Nevertheless, deeper knowledge into the molecular alterations by chronic opioid use in the human DLPFC and NAc is extremely limited.

We aimed to establish a more comprehensive understanding of the molecular changes across DLPFC and NAc in brains from subjects who were chronic opioid users also diagnosed with OUD. We used multiple levels of analysis by integrating transcriptomics across brain regions with traits related to OUD vulnerability using GWAS. Between the DLPFC and NAc, we found remarkable overlap in both upregulated and downregulated transcripts. Further investigation into these overlapping transcripts revealed pathways enriched for factors that control the formation and degradation of the extracellular matrix (ECM) and pro-inflammatory cytokine-mediated signaling. These pathways implicate neuroinflammation as a driver of ECM remodeling and synaptic reorganization, processes which are critical for opioid-induced neuroplasticity (14). Our analyses showed that microglia were central to these OUD effects in both the DLPFC and NAc. Finally, we found links between neuroinflammation and OUD, and strong associations with a genetic liability to risky behavior using GWAS. Our findings revealed novel genetic and molecular changes that may ultimately contribute to opioid dependence.

## Materials and Methods

Detailed procedures are provided in **Supplementary Methods.**

### Human Subjects

Brains were obtained during routine autopsies conducted by the Office of the Allegheny County of the Medical Examiner (Pittsburgh, PA) after consent was obtained from next-of-kin. Procedures were approved by the University of Pittsburgh’s Committee for Oversight of Research and Clinical Training Involving Decedents and Institutional Review Board for Biomedical Research. Each subject meeting diagnostic criteria for OUD at time of death (n=20) was matched with one unaffected comparison subject (n=20) for sex and as closely as possible for age (**Table 1; Table S1)**. Cohorts only differed by race (p=0.02). DLPFC (area 9) and NAc were identified on fresh-frozen right hemisphere coronal tissue blocks using anatomical landmarks (15), and tissue (∼50mg) was collected via cryostat, using an approach that minimizes contamination from white matter and other striatal subregions and ensures RNA preservation (16, 17).

**Table 1.** Subject summary demographic and tissue characteristics

| **Characteristic** | **Unaffected**<br>**comparison**<br>**(n=20)** | **Opioid**<br>**dependent**<br>**(n=20)** |
| --- | --- | --- |
| Age | 47.3 ± 9.5 | 46.9 ± 7.3 |
| Sex | 10M 10F | 10M 10F |
| Race | 13W 7B | 19W 1B |
| PMI (hours) | 15.7 ± 6.1 | 16.0 ± 5.3 |
| Brain pH | 6.6 ± 0.3 | 6.4 ± 0.2 |
| RIN | 8.0 ± 0.7 | 7.8 ± 0.7 |
| Tissue<br>Storage Time<br>(months) | 100.3 ± 86.1 | 103.0 ± 59.7 |

Values are mean ± SD. M, male; F, female; B, black; W, white

### RNA Sequencing analyses

Differential expression (DE) was assessed using limma with covariate selection (18). Transcripts with corrected p<0.01 and log_2_FC>±0.26 were considered DE (20–22). The top 250 DE transcripts were ordered by log_2_FC for unsupervised clustering of subjects. Biotypes of transcripts were determined using metaseqR (v3.11) (23). Overrepresentation of pathways (GO, KEGG, Hallmark, Canonical Pathways, Reactome, BioCarta, CORUM) was assessed using Metascape (http://www.metascape.org), with expressed transcripts as background. Networks were visualized with Cytoscape. INGENUITY**®** Pathway Analysis (Qiagen) and HOMER (v4.11) (24) was used to predict upstream regulators of DE transcripts. Rank-rank hypergeometric overlap (RRHO) (25, 26) was used to assess overlap of DE transcripts (p<0.01 in both regions).

### Identification of OUD-specific co-expression networks

We used weighted gene co-expression network analysis (WGCNA) to identify gene modules across samples (27, 28). Module differential connectivity (MDC) was used to quantify differences in co-expression within modules between OUD and unaffected comparison subjects. We used Fisher’s exact test to determine whether DE transcripts were enriched within WGCNA modules. ARACNe was used to identify hub and OUD-specific hub genes for network analysis (29) and Cytoscape was used to visualize networks. Overrepresentation of pathway categories for each module was assessed using Metascape, with the 5000 WGNCA-analyzed genes as background.

### Cell-type-specific DE analysis

We estimated cell-type fractions from bulk RNA-seq using Digital Sorting Algorithm (30) through deconvolution in BRETIGEA into astrocytes, endothelial cells, microglia, neurons, and oligodendrocytes (31). OUD and unaffected comparison subjects are compared for enrichment of each cell type using hypergeometric t-tests adjusting for brain region. We conducted cell-type-specific DE analysis with CellDMC (32) (FDR<0.05).

### Integration of DE transcripts with GWAS

Region-specific differentially up- and down-regulated transcripts (corrected p<0.01) were used to construct foregrounds for GWAS enrichment. We computed the partitioned heritability (GWAS enrichment) of brain region-specific noncoding regions containing and surrounding OUD transcript sets using the LD score regression pipeline for cell type-specific enrichment (33, 34). LD score regression coefficients were adjusted for FDR<0.1 on enrichments performed on all GWAS for foregrounds. A significant p-value indicates enrichment of the foreground genomic regions for GWAS SNPs relative to the background.

## Results

### Enrichment of DE transcripts involved in neuroinflammation and ECM remodeling in DLPFC and NAc in OUD

We first determined whether there were transcriptional differences by brain region in unaffected comparison subjects. Overall, the DLPFC and NAc had unique transcriptional profiles (**Fig.S2**). In DLPFC, many of the transcripts were related to synaptic vesicle transport (*e.g., KIF5C* (35, 36)*, STXBP5L* (37)), exocytosis (*e.g., STX1A* (38)), and neurotransmitter release (*e.g,. CADPS2* (39), *RIMS3* (40)) (**Data file S1**), while the NAc was enriched for cytoskeletal remodeling and chemotaxis (*e.g, ROBO1* (*41*)*, MTUS1* (*42*)) (**Data file S1**).

We next investigated the impact of OUD on region-specific transcriptional differences. Our results showed that opioid dependence had a profound effect on gene expression in DLPFC and NAc, with high numbers of DE transcripts in both brain regions (567 in DLPFC (**Data file S2**); 1306 in NAc (**Data file S3**; **Table S2**). The volcano plot of DE results for the DLPFC revealed an even distribution of downregulated (339) or upregulated (228) transcripts in OUD (**Table S2**, **Fig.1A**, **data file S2**). Unaffected comparison subjects were differentiated from OUD subjects by unsupervised clustering based on the expression of the top 250 DE transcripts (log_2_FC) in DLPFC (**Fig.1B**). The volcano plot for NAc revealed that many of the DE transcripts were downregulated (1085 compared to 221 upregulated transcripts) in OUD subjects (**Fig.1C**, **Data file S3**). Unsupervised clustering of subjects based on the top 250 DE transcripts in NAc identified subgroups in both the unaffected comparison and OUD cohorts (**Fig.1D**). Further analysis using principal components analysis (PCA) identified five subjects with OUD that clustered together with four unaffected subjects (**Fig.S3**). Each subject was diagnosed with an inflammatory disease (*e.g.,* asthma or arthritis; **Table S1**). Another three subjects from the unaffected comparison cohort also had histories of inflammatory disease, although these subjects seemed to cluster with the remainder of comparison cohort (**Fig.S3**). Overall, our findings suggest inflammatory disease may impact the NAc transcriptome, and further supports enrichment of neuroinflammation in subjects with OUD are independent of acute or chronic inflammation.

**Fig. 1.**
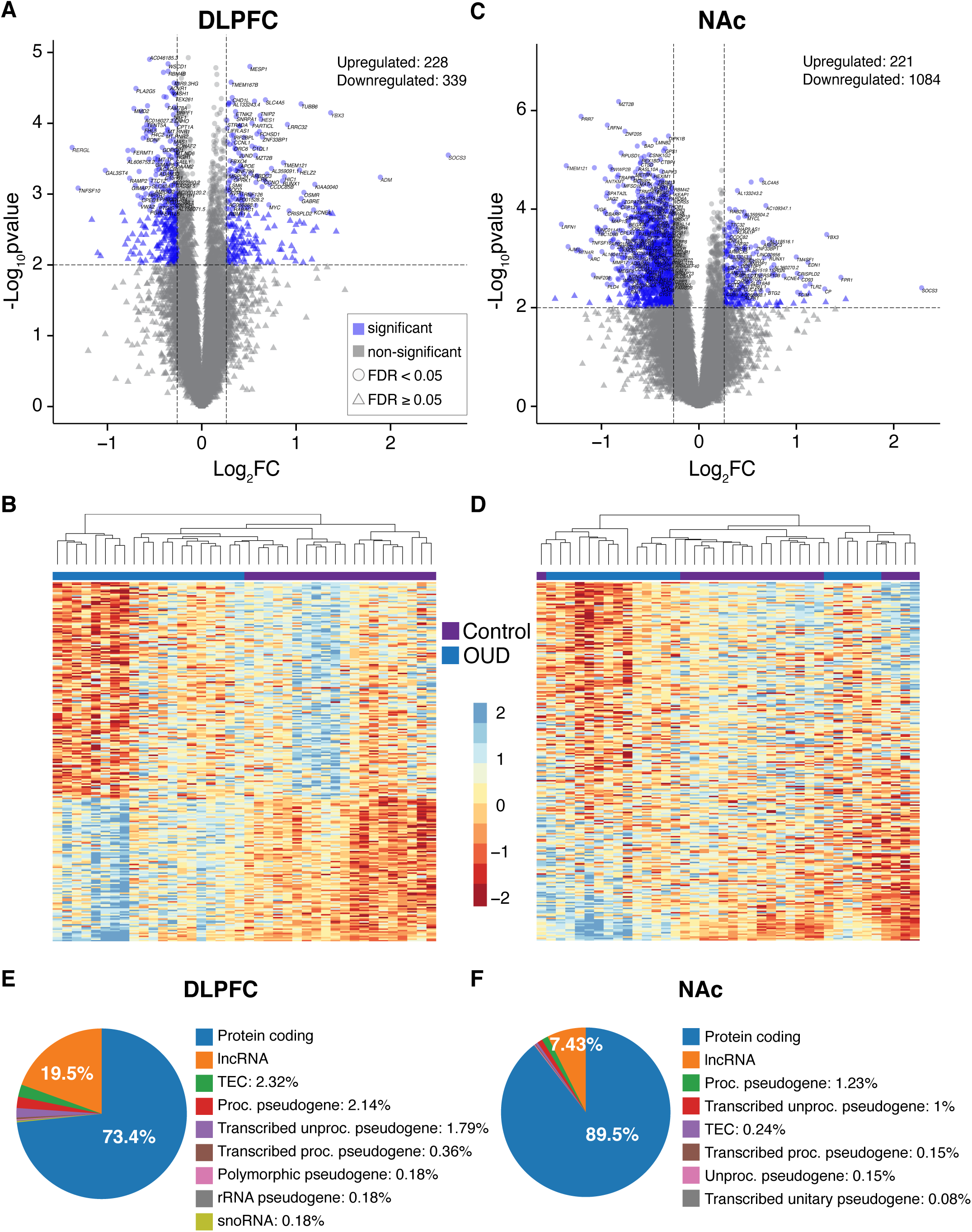
Transcriptomic changes in DLPFC and NAc from OUD subjects. **A.** Log_2_FC plotted relative to -log_10_p-value by volcano plot for DE genes in DLFPC. Horizontal dashed lines represent p-value significance cutoff of corrected p<0.01, while vertical dashed lines represent log_2_FC cutoffs of ≤-0.26 or ≥0.26 (FC≥1.2). Red circles represent DE genes that reach significance, log_2_FC, and FDR<0.05 cutoffs. **B.** Heatmap of the top 250 DE genes (corrected p<0.01 and log_2_FC of ≤-0.26 or ≥0.26) clustered by gene and subject. Each column represents a subject (unaffected comparison, purple; OUD, blue). Subjects within groups cluster together. **C.** Volcano plot for DE genes in NAc. Note the large numbers of genes that are significantly reduced in expression compared to genes that are increased. **D.** Heatmap of the top 250 DE genes clustered by gene and subject. Overall, subjects within groups cluster together with several groups of subjects forming separate clusters. **E.** Biotypes of DE genes (corrected p<0.01 and log_2_FC of ≤-0.26 or ≥0.26) in DLPFC. Protein coding genes represent the majority of DE genes (blue; 73.4%) followed by lncRNAs (orange; 19.5%). **F.** Biotypes of DE genes in NAc. Protein coding genes represent the majority of DE genes (blue; 89.5%) followed by lncRNAs (orange; 7.43%).

Most transcripts in both DLPFC and NAc were protein-coding and long noncoding RNAs (lncRNAs) (**Fig.1E-F**; DLPFC: 73.4% protein-coding, 19.5% lncRNAs; NAc: 89.5% protein coding, 7.43% lncRNAs). Further analysis revealed pathways associated with inflammation in both DLPFC and NAc (**Supplementary Results**, **Fig.S4-S5**). Other pathways included chondroitin/dermatan sulfate metabolism and synapse organization in NAc, suggesting links between extracellular matrix (ECM) remodeling, microglial cell migration, and synaptic plasticity, in line with recent work (43).

### High transcriptional coherence between DLPFC and NAc converges on neuroinflammatory and ECM pathways in OUD

Since pathways were largely similar between brain regions in OUD, we explored the extent of transcriptional overlap between DLPFC and NAc using rank-rank hypergeometric ordering (RRHO) (44, 45). RRHO orders transcripts in each brain region by both effect size direction and p-value. Substantial transcriptional overlap in both upregulated and downregulated transcripts was found between DLPFC and NAc (**Fig.2A**-**C**, Fisher’s exact test, p<10^-6^; **Fig.S6**). Such analyses may provide critical insight into the functional alterations across DLPFC—NAc circuits (3, 14, 46–49). Given the extent of transcriptional coherence, pathway enrichment analysis was conducted on DE transcripts shared between regions. Across DLPFC and NAc, the top shared pathways included those related to ECM (*e.g.,* biosynthesis of glycosaminoglycans and chondroitin sulfate) and inflammation (*e.g.,* cytokine-mediated immune signaling via tumor necrosis factor alpha (TNFα) and nuclear factor kappa B (NFκB) (**Fig.2D**). Other pathways were associated with cellular stress responses (*e.g.,* DNA damage repair), and protein and histone modifications (*e.g.,* ubiquitination and acetylation) that govern epigenetic regulation (**Fig.2D**, **Fig.S7**).

**Fig. 2.**
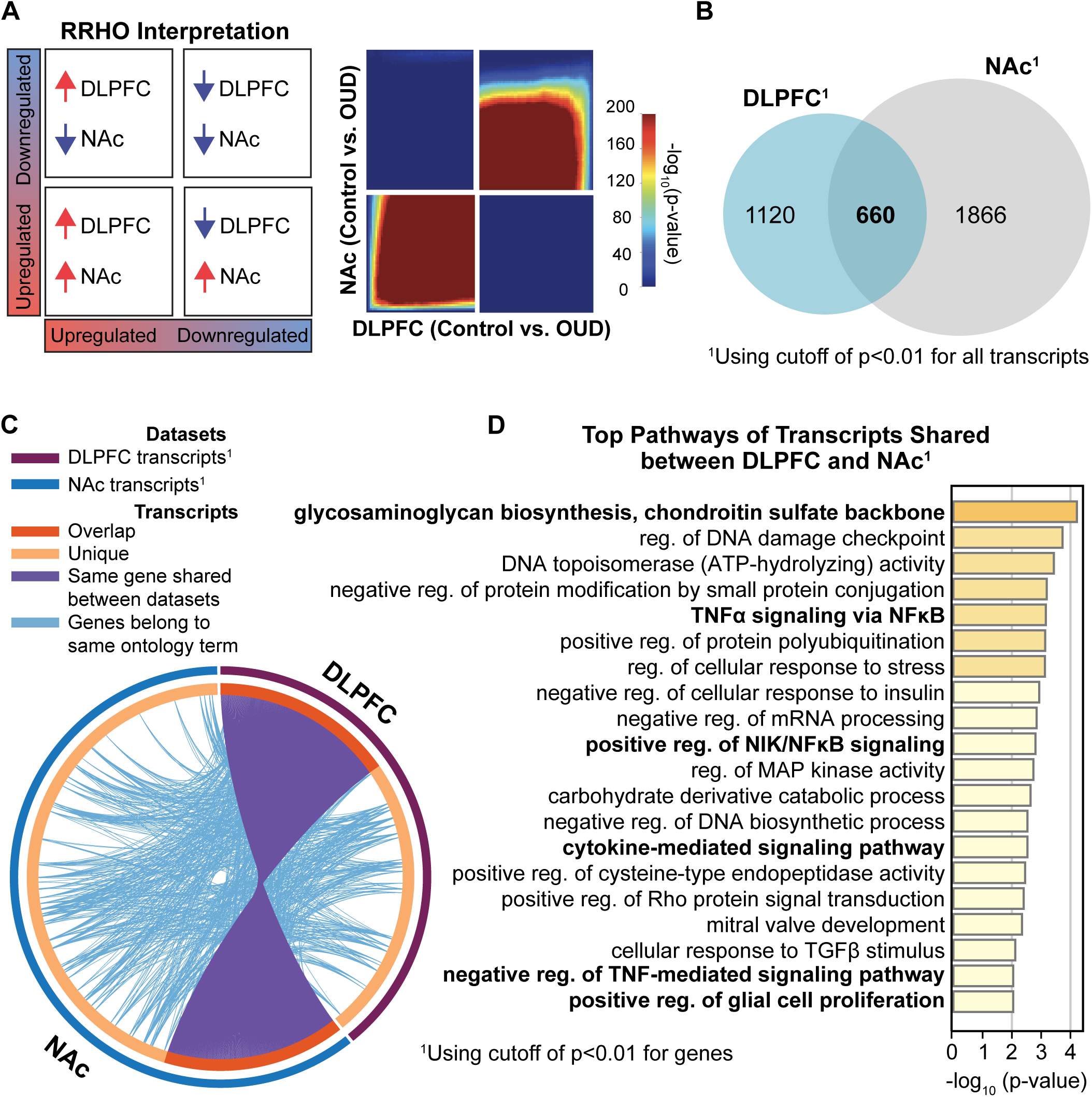
High transcriptional concordance reveals commonly altered molecular pathways between brain regions associated with opioid dependence. **A.** RRHO plot indicating high degree of overlap, or transcriptional concordance, between DLPFC and NAc in OUD subjects. **B.** Venn diagram of DE genes between DLPFC and NAc. **C.** DE genes (purple lines) and their ontology (light blue lines) highly overlapped (orange) between DLPFC and NAc. **D.** Top 20 pathways of DE genes shared between DLPFC and NAc. Many of these pathways are related to ECM and neuroinflammatory signaling.

Given the association with epigenetic regulation, we investigated the enrichment of chromatin state by comparing the transcription start sites from DE transcripts in OUD to states previously defined in postmortem brains from unaffected comparison subjects using genome-wide maps of epigenetic modifications (50). Transcription start sites of DE transcripts in DLPFC were enriched for genomic regions marked by weak polycomb repression (**Fig.S8A**, p<10^-5^). Weak polycomb repression diminishes histone marks (*e.g.,* H3K27me3), which would otherwise inhibit transcription at the start site (50). Relative to unaffected subjects, the enrichment of upregulated DE transcripts in DLPFC from OUD subjects suggests that opioids induce the activation of otherwise repressed areas of the genome to promote transcription (Figure S6). In NAc, DE transcripts were enriched for genomic regions marked for a quiescent state (**Fig.S8A**; p<10^-6^). Quiescent states are characterized by the complete absence of histone marks linked to transcriptional inactivity (50). Similar to DLPFC, such findings suggest opioids activate otherwise inactive genomic regions, resulting in upregulation of specific transcripts in NAc (**Fig.S8B**). Overall, these findings reflect a fundamental disturbance of chromatin states and transcriptional regulation by opioids relative to baseline epigenetic states.

### Upregulation of microglia markers in OUD

By leveraging the high transcriptional coherence between DLPFC and NAc, we identified cell types with specific DE transcripts expressed in both brain regions using deconvolution (32). Transcripts were clustered in major brain cell types including astrocytes, endothelial cells, microglia, neurons, and oligodendrocytes and oligodendrocyte precursor cells. Compared to unaffected subjects, markers for microglia (*e.g.*, *ZNF812*) were significantly enriched in OUD (p=0.03; **Fig.3A**). We next conducted cell type-specific DE analysis in DLPFC and NAc by identifying DE transcripts that significantly co-varied with specific cell markers. Using brain region as a covariate, we detected a single DE transcript associated with microglia markers, *HSD17B14* (**Table S3**). Previous work showed that reduced expression of *HSD17B14* in microglia resulted in exaggerated pro-inflammatory glial activation (51). This further supports evidence for neuroinflammatory processes in DLPFC and NAc in OUD.

**Fig. 3.**
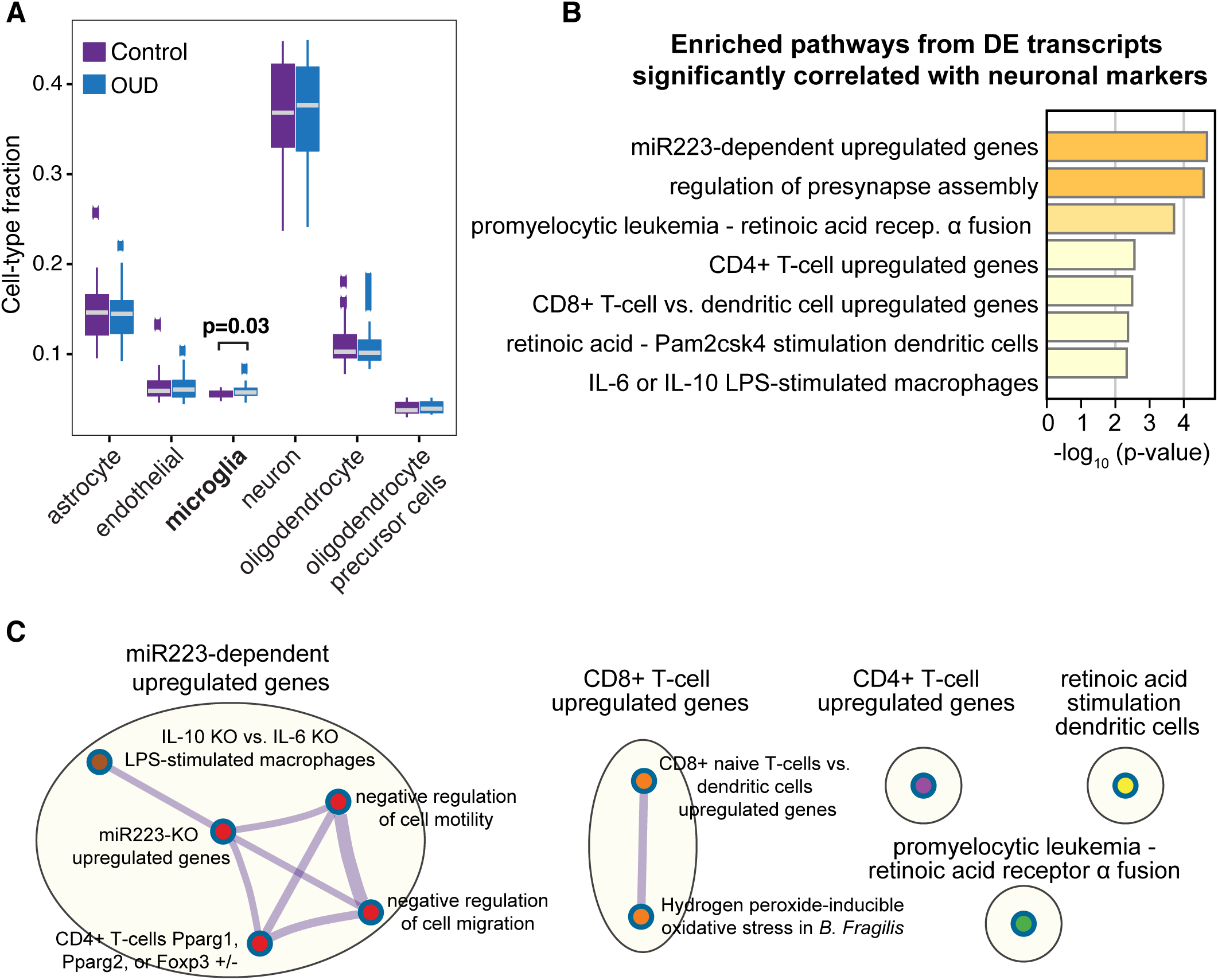
Predicted microglial cell-type enrichment in both DLPFC and NAc. **A.** Deconvolution analyses indicates modest enrichment of microglia gene markers in DLPFC and NAc of OUD subjects (p<0.03; hypergeometric t-test controlling for brain region), consistent with enrichment of pathways in DE genes related to immune function. **B.** Pathway enrichment analysis on the significantly altered DE genes in neurons identified pathways related to inflammation**. C.** Significantly enriched terms based on pathways included in multiple annotated sets (*e.g.,* GO, KEGG, hallmark, etc.) using hypergeometric p-values and enrichment factors. The network is visualized with Cytoscape using Community cluster with categorical labels representing multiple clusters. Individual nodes are also labeled. Note nodes are related to presynaptic structure and function and miR233-dependent regulation of macrophage and T-cell activation.

Focusing on transcripts that correlated with changes in neuronal markers, we identified 25 DE transcripts significantly enriched for pathways related to neuroimmune signaling between neurons and local inflammatory cells (*e.g.,* microglia) (**Fig.3B**, **Table S3**). We found several upregulated genes that are controlled by a single microRNA (miR), miR223 (**Fig.3B**). miR223 is a crucial modulator of macrophage activation (52), further implicating microglia in the regulation of neuroimmune responses in OUD. A broader view indicated that immune-related pathways were largely distinct between microglia and other immune cell types including CD4^+^ and CD8^+^ T-cells (**Fig.3C**). This highlights the possibility of interactions between neurons and other immune cells in brains of OUD subjects. Lastly, we also found strong enrichment of DE transcripts with synapse-related pathways (**Fig.S9**).

### Increased connectivity of neuroinflammatory and ECM signaling gene modules in OUD

We used WGCNA to investigate correlations among transcripts in both unaffected comparison and OUD subjects. WGCNA identified 17 co-expression modules in DLPFC and 15 modules in NAc (**Fig.4A**). To identify OUD-specific modules, we used MDC to directly compare network connectivity of each module between OUD and unaffected subjects. More coordinated expression of transcripts in unaffected subjects relative to OUD subjects indicates a ‘gain’ of connectivity in unaffected subjects. Conversely, a module that loses connectivity in unaffected subjects is more coordinated in OUD. In DLPFC, 15/17 modules lost connectivity and two remained unchanged in unaffected subjects (**Fig.4B**). In NAc, 13/15 modules lost connectivity, while one module gained connectivity, and another remained unchanged in unaffected subjects (**Fig.4B**). These findings indicate that overall module structures were largely distinct between OUD and unaffected comparison subjects.

**Fig. 4.**
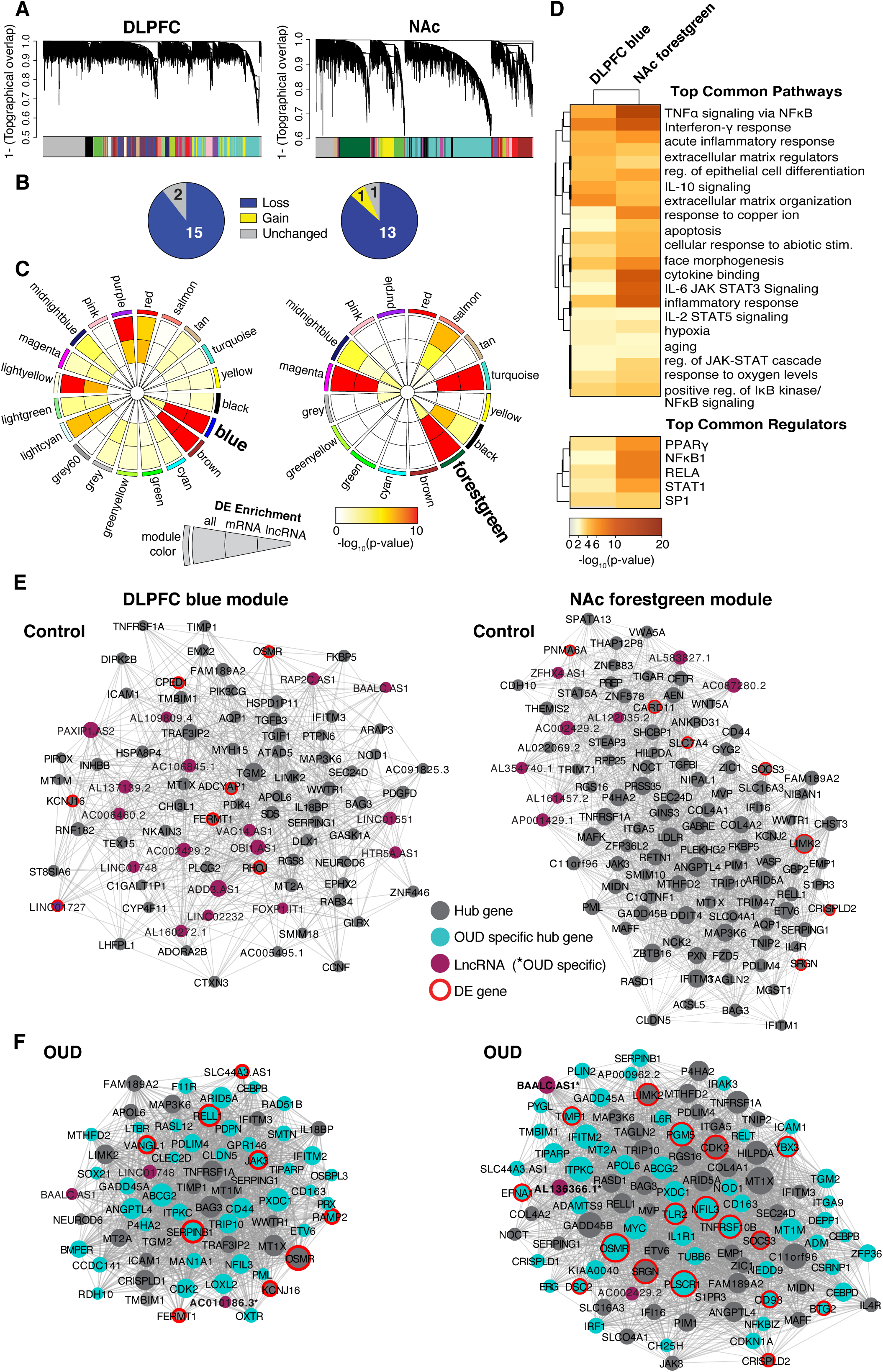
OUD associated gene networks in DLPFC and NAc. **A.** Weighted gene co-expression network analysis (WGCNA) was used to generate co-expression modules, with the network structure generated on each brain region separately. The identified modules that survived module preservation analysis were arbitrarily assigned colors and the dendrogram shows average linkage hierarchical clustering of genes. **B.** Pie charts summarize results from the module differential connectivity (MDC) analysis compared OUD to unaffected comparison subjects. The majority of modules were lost in unaffected comparison subjects, revealing OUD-specific modules in both DLPFC and NAc. **C.** Circos plot identified by module names and colors. Enrichment for full list of differentially expressed (DE) genes, protein-coding (mRNA), and long non-coding RNAs (lncRNA), are indicated by semi-circle colors within each module, with increasing warm colors indicating increasing-log_10_ p-value. LncRNA enrichment was examined based on the high prevalence of these transcripts in the Biotype analysis (see Figure 2a,e). MDC analysis indicated a loss of connectivity in the DLPFC blue module and NAc forestgreen module. These modules were also enriched for DE mRNA and lncRNAs. **D.** Pathway enrichment analysis compared gene networks within the DLPFC blue module and NAc forestgreen module. Warmer colors indicate increasing-log_10_ p-value and highly shared pathways between the modules. Hub gene co-expression networks of the DLPFC blue module **E.** unaffected comparison subjects and **F.** OUD subjects, and networks of the forestgreen module in NAc **E** unaffected comparison subjects and **F.** OUD subjects. Node size indicates the degree of connectivity for that gene. Turquoise nodes indicate OUD-specific hub genes, purple nodes indicate LncRNA gene, gray nodes indicate hub genes, and red halos indicate DE genes. Edges indicate significant co-expression between two particular genes.

To further investigate the biological significance of OUD-specific modules, we examined DE enrichment within each module focused on highly represented mRNAs and lncRNAs (**Fig.1E-F**). The DLPFC blue module and the NAc forestgreen module were of particular interest because of their increased connectivity in OUD compared to the other modules and significant enrichment of both DE mRNA and lncRNA transcripts in OUD (DLPFC blue full DE list q<10^-26^, mRNA q<10^-23^, lncRNA p=0.002; NAc forestgreen full DE list q<10^-23^, mRNA q<10^-22^, lncRNA p=0.004). The topography of DLPFC blue and NAc forestgreen networks were more correlated in OUD relative to unaffected comparison subjects (**Fig.4E-F**), consistent with strengthening of module connectivity.

Given the substantial overlap in transcripts and pathways across DLPFC and NAc, we next tested the degree of overlap in transcript co-expression networks between DLPFC blue and NAc forestgreen modules. Indeed, there was substantial overlap in transcripts between these modules (Fisher’s exact test, p<10^-16^). These modules were enriched for pathways related to neuroinflammation and ECM remodeling, consistent with our above findings (**Fig.4D, Fig.S10**). The top shared pathways include TNFα signaling via NFκB, interferon-γ response, and acute inflammatory response via IL-6 (53) and IL-2 (54) signaling (**Fig.4D**). Each of the top shared upstream regulators are modulators of inflammatory response: PPARγ (55), NFKB1 (56), RELA (56), STAT1 (57), and SP1 (58) (**Fig.4D**). Notably, we also found pathways related to ECM remodeling (**Fig.4D**). These data build upon our above analyses highlighting critical roles for neuroinflammation and ECM in OUD.

To identify potential drivers of co-expression networks, we detected highly connected “hub” genes within a module that were predicted to regulate the expression of other module genes. Many of the hub genes that were specific to OUD in DLPFC and NAc included inflammatory regulators, such as *JAK3* (59), *SERPINB1* (60), and *RELL1* (61) in DLPFC (**Fig.4F**), and *TLR2* (62), *TNFRSF10B* (63), and *NFIL3* (*64*) in NAc (**Fig.4F**). We also found additional classes of highly connected transcriptional regulators within our co-expression networks including lncRNAs and RNA-binding proteins. Several lncRNAs were highly connected in OUD-specific networks in DLPFC and NAc (*e.g.,* DLPFC: *AC0101086.3* and NAc: *BAALC.AS1*) (**Fig.4F**, **Table S4**). *AC0101086.3* is located proximal to *CLEC2D* in the genome, another OUD-specific hub gene. *CLEC2D* encodes ELT1, an activator of innate immunity (65, 66). *BAALC.AS1* is highly expressed in brain and regulates astrocytes (67). In the NAc forestgreen module, we also identified *YBX3*, an RNA-binding protein. YBX3 regulates the expression of large neutral amino acid transporter 1 (LAT1) (68), which is necessary for the uptake of catecholamine precursor, L-DOPA in dopaminergic cells (69). Notably, changes in *YBX3* expression in human midbrain has previously been implicated in opioid dependence (70). We identify putative mechanisms involved in neuroinflammation and dopamine neurotransmission in OUD.

### Associations between DE transcripts in NAc and genetic liability for substance use-related traits in OUD

Opioid dependence is strongly linked to impulsivity and risk-taking. Given our results highlighting inflammation in OUD, we determined whether these pathways were linked to traits related to opioid dependence. To test this, we employed a GWAS-based approach that integrated risk loci associated with substance use-related traits (*e.g.,* opioid dependence, smoking, and risky behavior) and psychiatric disorders (71–73) with transcriptional profiles in DLPFC and NAc from OUD subjects. Loci identified by GWAS are known to overlap with intronic and distal intergenic noncoding regions within *cis*-acting regulators of gene expression (50). Using this information, we examined whether noncoding regions proximal to our DE transcripts were enriched for genetic risk variants associated with vulnerability to opioid dependence. We discovered significant enrichment of downregulated DE transcripts in NAc of OUD subjects for genes associated with smoking initiation and cessation (**Fig.5A**), along with attention deficit hyperactivity disorder (ADHD), bipolar disorder, depression, and risky behavior GWAS (**Fig.5C**, FDR<0.05). Many of these transcripts were associated with neuroimmune signaling, along with the machinery involved in synaptic neurotransmission (**Fig.2**, **Fig.4**). There was no enrichment of DE transcripts in chronic pain, opioid dependence, or unrelated GWAS traits (*i.e.,* bone mineral density, coronary artery disease, and lean body mass) (**Fig.5A-D**). Our results therefore bridge substance use related genetic risk factors to our transcriptomic findings in OUD.

**Fig. 5.**
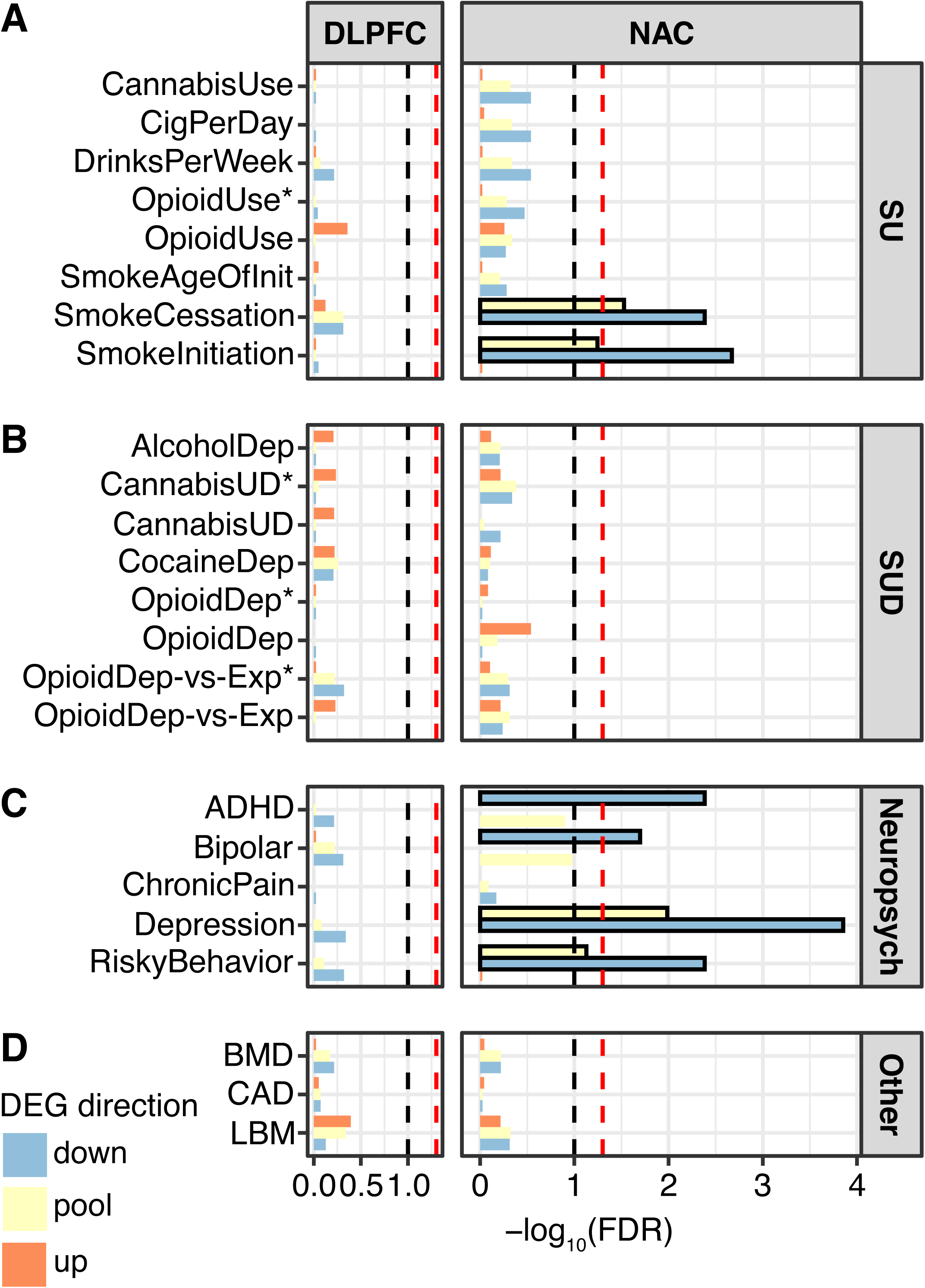
Differentially expressed genes in the DLPFC and NAc enrich for genetic liability of risky behavior. Several well-powered genome-wide associated studies (GWAS) have identified risk loci associated with substance use (SU), substance use disorder (SUD)-, and neuropsychiatry (neuropsych)-related traits. Significant risk loci overlap with intronic and distal intergenic noncoding regions, presumably within cis-acting regulatory elements of gene expression. **A.** Proximal noncoding regions of differentially expressed (DE) genes in DLPFC and NAc from OUD subjects were investigated for enrichment of genetic risk variants of SU-related traits using partitioned heritability linkage-disequilibrium score regression analysis. No enrichment was found in the DLPFC, but significant enrichment was found in the NAc for up and down-regulated genes with smoking cessation and smoking initiation. **B.** No enrichment was found in the DLPFC or NAc for SUD-related GWAS. **C.** No enrichment was found for neuropsychiatry-related GWAS in the DLPFC, but there was enrichment in the NAc for attention deficit hyperactivity disorder (ADHD; down-regulated), bipolar disorder (downregulated), depression (up and downregulated), and risky behavior (up and downregulated) GWAS. **D.** No enrichment in DLPFC or NAc for unrelated GWAS traits, including bone mineral density (BMD), coronary artery disease (CAD), and lean body mass (LBM). *, indicates in African American population; lack of asterisk indicates European American population.

## Discussion

Our work demonstrates that inflammation is the most salient biological process identified in OUD. Clinical findings that link inflammation to OUD are primarily from reports of elevated levels of circulating pro-inflammatory cytokines in opioid dependent individuals (74). Pro-inflammatory cytokines in periphery and brain may have functional consequences that contribute to OUD. In line with this, microglial inhibition mitigates subjective withdrawal (75), reduces motivation to consume opioids in dependent individuals (76), and attenuates opioid conditioned reward and seeking in rodents (77–80). However, the unanswered question was whether these findings were clinically relevant for OUD. By focusing on key regions involved in OUD, we reveal several key inflammatory pathways altered in OUD in DLPFC and NAc.

One of the top pathways shared between DLPFC and NAc is TNFα signaling via NFκB. Receptors known to activate NFκB, TNF and TLR4 (81, 82), are among the top predicted upstream regulators of DE transcripts. Though still controversial, there is growing evidence that opioids can also induce a neuroinflammatory response via direct activation of TLR4, a transmembrane receptor which activates NFκB signaling and inflammatory cascades (83). Upon activation, NFκB dimers translocate to the nucleus to drive transcription of cytokines, chemokines, and interleukins (81, 82). Additionally, opioids can activate NFκB via opioid receptors (85–89), and activation of NFκB signaling can, in turn, promotes transcription of opioid receptors and peptides (90–95), involved in opioid reward (90, 96, 97). Thus, while opioids can influence immune function via NFκB, opioids may also activate NFκB, with downstream effects on addiction-related behaviors, independent of immune function.

An effective neuroinflammatory response involves the interplay between immune cells and local ECM remodeling. The ECM is an assembly of adhesion molecules, proteins, polysaccharides, and proteoglycans, critical for BBB integrity and synaptic function (98). Both the formation and degradation of ECM in the brain depend on the aggregation of a specific family of proteoglycans, chondroitin sulfate glycoaminoglycans (CS-GAGs) (100). Significantly, CS-GAGs constitute the top shared pathway enriched between DLPFC and NAc in OUD subjects. CS-GAGs aggregate in the perisynaptic space in response to inflammation. Increasingly, ECM remodeling is also implicated in synaptic plasticity (101), with roles in neurite outgrowth, dendritic spine formation and morphology (100, 102), and myelination (103, 104). Indeed, reduced myelination has been found in clinical neuroimaging and postmortem brain studies of chronic opioid users (105–109), as well as in rodent models of chronic opioid exposure (110, 111). Importantly, the cytokines we identified in DLPFC and NAc of OUD subjects, such as interferons and TNFs, modulate ECM remodeling (112, 113), and therefore, directly link neuroinflammation, ECM, and opioids.

We therefore posit that opioid-induced changes in CS-GAG signaling driven by neuroinflammation disrupt ECM structure and have profound consequences on dendritic, synaptic, and behavioral plasticity. For example, reorganization of ECM in NAc via matrix metalloproteinases (MMPs), which facilitate matrix degradation and reassembly, leads to increased potentiation of glutamatergic synapses (114) and opioid relapse (115). Importantly, we identified *TIMP1* and *TLR2* as OUD-specific hub genes, both of which have crucial roles in ECM remodeling (116) and functional reorganization of excitatory synapses by directly inhibiting MMPs (114). We therefore speculate: 1) opioid use elicits release of pro-inflammatory cytokines in the brain that activate TIMP1 and TLR2, protein modulators of ECM organization (117); 2) these activated modulators modify MMP activity to alter ECM organization; and 3) these disturbances to ECM remodeling alter synaptic plasticity^136^ and result in the behavioral changes related to OUD. Future experiments using animal models will directly test these possibilities.

Multiple lines of evidence point to the centrality of microglia, the primary resident immune cells in the brain, in the pathways we have identified in OUD subjects. First, microglial markers were significantly enriched across DLPFC and NAc in OUD. Moreover, our cell-type marker-based approach revealed that the top DE transcript, *HSD17B14*, was highly correlated with microglial markers. *HSD17B14* maintains microglial homeostasis during inflammation (51). Second, our cell-type marker analysis revealed pathways involved in microglial activation and the activation of additional immune cell types, including CD4^+^ and CD8^+^ T-cells. Earlier work suggests synergistic relationships between microglia and T-cells that may potentiate neuroinflammation in OUD (118). Third, the cytokines we identified, including IL-1β, IL-6, IL-2, and TNFα are secreted by microglia (119), all of which are implicated in opioid reward and dependence (120–122). Fourth, pathways identified in OUD map are related to transcriptional regulation in microglia during inflammation. Specifically, network analyses predicted the following top upstream regulators as PPARγ, NFκΒ1, RELA, STAT1, and SP1. While microglia rely on multiple families of transcription factors (123), our findings in OUD largely identify transcription factors that are important for guiding microglial response to inflammatory signals. For example, PPARγ is a nuclear transcription factor preferentially expressed in human microglia. PPARγ is activated in response to neuroinflammation and leads to anti-inflammatory responses that are neuroprotective (123). Aberrant activation of STAT1 in microglia upregulates several pro-inflammatory cytokines. Intriguingly, STAT1-dependent signaling regulates the expression of the human μ-opioid receptor via IL-6, another pro-inflammatory cytokine that we identified in our analyses (124). Finally, we found two pathways intimately linked to microglial function: ameboidal migration and integrin signaling. Integrins physically tether microglia and neurons to the ECM scaffold (125). Microglia rely on the ECM for ameboidal migration, further linking microglia, ECM, and neuroinflammation in OUD.

In addition to changes in protein-coding transcripts, we discovered that subjects with OUD exhibited marked expression changes in lncRNAs. LncRNAs are key regulators of gene and protein expression (126). Several of the lncRNAs we identified in OUD subjects, *AC0101086.3, BAALC.AS1,* and *AL136366.1*, are implicated in both neurotransmission and neuroinflammation. In DLPFC, we found *AC0101086.3* to be an OUD-specific hub lncRNA. This lncRNA is located proximally to *CLEC2D*, which encodes ELT1, the functional ligand for the natural-killer cell receptor NKR-P1A. NKR-P1A controls immunosurveillance via natural-killer, dendritic, and B-cells (65, 66). In NAc, we found *BAALC.AS1* (127) and *AL136366.1* (128) as OUD-specific hub lncRNAs, both of which are involved in various brain functions (129, 130).

Functional alterations occur at the circuit-level across multiple brain regions in opioid dependence and treatment (131). One such change occurs at the transcriptional level, where different brain regions can increasingly synchronize their patterns of transcription. There is evidence that increases in transcriptional synchrony across brain regions occur in response to insults including stress and drugs of abuse (20, 132). Consistent with this, we found significant transcriptional synchrony between DLPFC and NAc in OUD. To date, the relevance of such synchrony remains unclear. We speculate that, because the ranking, effect size, and direction of change in numerous transcripts is highly similar between brain regions, such synchrony may represent a common pathophysiological response to neuroinflammation in OUD. Clearly, further work is required to explore these possibilities.

Integrating large-scale gene expression profiles with relevant GWAS findings reveals novel gene-trait associations in OUD. We demonstrated that downregulated genes in the NAc of OUD subjects were significantly enriched for GWAS of risky behavior (71). Risky behavior forms a functional triad with mood and impulsivity (133), where impulsivity is a risk factor for substance use (134–136). Our findings therefore support relationships between genetic risk, brain region-specific transcriptional changes, and vulnerability to OUD.

Overall, our data reveal new connections between the brain’s immune system and opioid dependence in the human brain. These results provide a novel putative mechanism across transcriptional networks, biological pathways, and specific cell types for the detrimental neuroadaptations across corticostriatal circuitry that result from chronic opioids and OUD. The insights provided by our work offer the opportunity for new therapeutic targets with improved efficacy to treat OUD.

## Acknowledgments

We would like to thank the staff and technicians who work diligently as part of the Brain Tissue Program at the University of Pittsburgh. Human tissue was obtained from the NIH NeuroBioBank and the University of Pittsburgh Brain Tissue Donation Program. This study was funded by the Hamilton Family Prize for Basic Neuroscience Research in Psychiatry at the University of Pittsburgh School of Medicine to R.W.L., NHLBI R01HL150432 to R.W.L, and NIDA R01DA051390 to R.W.L and M.L.S.

## Author Contributions

M.L.S., J.R.G., D.A.L., G.C.T. and R.W.L. designed and coordinated the study. X.X., W.Z., J.W., B.N.P., C.S., A.R.P., Z.F., G.C.T., and R.W.L. conducted statistical analysis and data interpretation. S.M.K., M.A.H., J.R.G., and M.A.S. obtained samples and processed samples for RNA-sequencing. M.L.S. and R.W.L. obtained funding for the project, and M.L.S., Z.F., and R.W.L. drafted the manuscript.

