## Supplement for "Transcriptional alterations in opioid use disorder reveal an interplay between neuroinflammation and synaptic remodeling"

617-358-9563

**Supplementary Materials and Methods**

*Human subjects*

An independent committee of clinicians made consensus, lifetime DSM-IV diagnoses for each subject using the results of a psychological autopsy, including structured interviews with family members and review of medical records, as well as toxicological and neuropathological reports (12). The same approach was used to confirm the absence of lifetime psychiatric and neurologic clinical disorders in the unaffected comparison subjects (controls). Subject groups did not differ in mean age, postmortem interval (PMI), RNA integrity number (RIN), or tissue storage time at ­80^o^C (all t_38_<1.18, all p>0.25). Brain pH significantly differed between diagnostic groups (t_38_=2.54, p=0.015); however, the mean difference was 0.2 pH units and of uncertain biological significance. Subject groups significantly differed in racial composition (χ^2^=5.63, p=0.018).

*RNA-sequencing and analyses*

Tissue was homogenized in TRIzol (Invitrogen, Carlsbad, CA) and RNA extracted using the RNeasy Lipid Tissue Mini Kit (Qiagen). RNA integrity was assessed using a Bioanalyzer (Agilent) and concentration was determined using a Qubit fluorometer (Invitrogen). Libraries were constructed using TruSeq Total RNA sample kit and sequenced by Illumina NextSeq 500. All samples were sequenced across all lanes to avoid lane bias by a technician blind to treatment groups (paired-end 75 cycles). Average sample coverage was 29 million reads before alignment. FastQC was used to assess the quality of the data. Per base sequence quality was high (mean Phred>30), indicating good quality of the experimental data. HISAT2 was used to align the read to the Ensembl human reference genome (GRCh38.p13) reference (16). The mean reads mapped was 97.52% +/- 0.002 standard deviation. Principal component analysis confirmed the absence of batch effects for each brain regions (**Fig.S1**). After alignment, Bam files were converted to expression count data using HTSeq (17). The RNAseq count data were transformed to log*2* count data using voom function of the Bioconductor limma package (18, 19). A total of 80 samples (20 control DLPFC, 20 control NAc, 20 OUD DLPFC, 20 OUD NAc) with 60,676 genes preprocessed together. Low expression genes were filtered out and 15,042 genes remained (retained genes with log_2_ transformed counts per million greater than 1 in 50% or more subjects). Briefly, for each gene, we first adjusted the difference of expression (log_2_FC) with covariates (sex, age, race, PMI, pH, RIN) by including each covariate in the regression model. Next, at most 2 covariates were selected to be included in the full model using Bayesian information criterion (BIC) for each gene individually (21). We used HOMER (v4.11) to predict upstream factors that might drive differential expression patterns (27). HOMER determines if binding motifs within a gene set are enriched compared to what is expected in the background gene list; all promoters (except those chosen for analysis) were used as background.

We used the threshold-free approach rank rank hypergeometric overlap (RRHO) (36, 37) to assess overlap of differential expression patterns between the DLPFC and NAc. RRHO identifies overlap between two ranked lists of differential gene expression. For each of the two datasets, RRHO ranks the entire gene list by DE p-value and effect size direction; one dataset is represented on the X-axis and one on the Y-axis.

*Chromatin states of DE genes*

Chromatin states from the 15-state model trained on histone modifications were obtained from the Roadmap Epigenomics Project (27). Genome state annotations were converted to hg38 from hg19 (28). ENSEMBL genes (v79) were matched to chromatin states based on the annotated transcription start site (29). Gene expression differences measured in the DLPFC in OUD subjects compared to control subjects were compared to PFC chromatin states. No chromatin states were available for the NAc, so chromatin states for the anterior caudate were used as a substitute, as both regions are dominated by medium spiny neurons. The hypergeometric test was used to calculate enrichment of observed overlaps in the OUD-higher and OUD-lower gene sets of both brain regions to each chromatin state in corresponding brain region.

*Weighted gene co-expression network analysis (WGCNA)*

WGCNA was performed using the 5000 most variable genes across samples. Modules were built using OUD samples and module preservation analysis was performed to determine module robustness (33). Module differential connectivity (MDC) was used to quantify differences in co-expression between OUD and controls. MDC is the ratio of the connectivity of all gene pairs in a module in OUD subjects to that of the same gene pairs in controls (34). MDC>1 indicates gain of connectivity, while MDC<1 indicates loss of connectivity. To statistically test the significance of MDC, we estimated the p-value based on two types of shuffling schemes: 1) shuffled samples (adjacent matrix with non-random nodes but random connections); 2) shuffled genes (adjacent matrix with random nodes but non-random connections). Data were permuted 1000 times. P-values were converted to q-values following Benjamini Hochberg procedure, and MDC q<0.05 was considered significant.

Overrepresentation of pathway categories (GO, KEGG, Hallmark, Canonical Pathways, Reactome, BioCarta, CORUM) for each module was assessed using Metascape (http://www.metascape.org), with the 5000 WGNCA-analyzed genes as background. To examine overlap of pathways, pathways are hierarchically clustered into a tree based on Kappa-statistical similarities among gene memberships (0.3 Kappa score used as threshold to cast tree into term clusters and thickness of edge represents similarity score). A subset of representative terms from this cluster were selected for network layout, where each term is represented by a circle node (size proportional to number of input genes within term and color intensity reflects p-value).

ARACNe was used to identify hub and OUD-specific hub genes for network analysis (35). A gene is considered a hub if the N-hob neighborhood nodes (NHNN) is significantly higher than the average NHNN. A hub gene is considered an OUD-specific hub if it attains hub status in OUD, but not control subjects. Cytoscape 3.8.0 was used to generate networks of hub genes; only correlations >0.98 are included in the networks.

*Relationship of identified DE lists to GWAS*

We selected intronic and intergenic regions of a marker gene from the GENCODE human hg38 comprehensive gene annotation release 33. For each set of gene markers, we took the region spanning the gene body and subtracted exons to define introns. Then, we took the 20kb regions flanking the gene body as the proximal intergenic regions. To avoid collision with nearby genes in dense, gene-rich loci, we removed flanking sequences that overlap exons. We then merged intronic and intergenic regions for each brain region and log_2_FC gene set foreground. We found that the LD score (LDSC) GWAS enrichment analysis is sensitive to selected background set of matched regions for proper enrichment above the foreground set. We combined intronic and flanking intergenic regions for all hg38 genes from the GENCODE annotation to create the background for LDSC GWAS enrichment analysis.

The summary statistics of GWAS SNPs used in these analyses were downloaded and processed using the LDSC munge_sumstats function to filter rare or poorly imputed SNPs. The HapMap SNPs excluding the MHC regions used to filter SNPs were downloaded from <http://ldsc.broadinstitute.org/static/media/w_hm3.noMHC.snplist.zip>. Column headings of each GWAS file were inspected to ensure the effect allele, non-effect allele, sample size, p-value, and signed summary statistic for each SNP in each GWAS were included and appropriate for the LDSC pipeline.

The LD scores were estimated for each foreground set and corresponding background set with the LD score regression pipeline make_annot and ldsc functions using hg38 1000 Genomes European Phase 3 (1000G EUR) cohort plink files downloaded from <https://data.broadinstitute.org/alkesgroup/LDSCORE/GRCh38/>. The baseline v1.2 files for cell type-specific enrichment in hg38 coordinates were downloaded from the same link above and the corresponding weights `weights.hm3_noMHC’ file excluding the MHC region from <https://data.broadinstitute.org/alkesgroup/LDSCORE/>. We find that HapMap SNPs and corresponding weights file used in the LDSC analyses only referred to rsIDs, rather than genomic coordinates, so only the baseline and LD statistics used to annotate the foreground and background files have to be in hg38 coordinates. An `enrichment.ldcts’ file listing the annotated foreground/background pair was created for each foreground. The partitioned heritability was estimated using the ldsc function, which integrates the foreground and background LD score estimates, munged GWAS SNP data, baseline variant data, and variants weights. The final function call to GWAS enrichment is as follows:

python ldsc.py --h2-cts $Munged_GWAS \

--ref-ld-chr baseline_v1.2/baseline. \

--w-ld-chr weights.hm3_noMHC. \

--ref-ld-chr-cts enrichment.ldcts \

--out $Output_Label

The traits of brain-related GWAS include multisite chronic pain(43), tendency to engage in risky behavior (44), schizophrenia risk (45), opioid dependence (46), cannabis use (47), diagnosis of obsessive-compulsive disorder (48), age of smoking initiation (49), heaviness of smoking (49), alcoholic drinks per week (49), having ever regularly smoked (49), current versus former smokers (49), highest level of educational attainment (50), sleep duration (51). The traits of the non-brain related traits include lean body mass (52), bone mineral density (53), and coronary artery disease (54). The pipeline produced LD score regression coefficient, coefficient error, and coefficient p-value estimates.

**Supplementary Results**

To determine the identity of the genes encoded by these transcripts, we focused on protein-coding and long noncoding RNAs (lncRNAs) because they were prominent in both brain regions in OUD subjects. We examined the functions of these DE transcripts via pathway analysis. The top pathways represented by DE transcripts in the DLPFC were tumor necrosis factor (TNFα) signaling via nuclear factor kappa B (NFκB), as well as adipocytokine and interferon gamma signaling, clearly implicating an inflammatory response in DLPFC in OUD (**Fig.S3A**). We also found pathways related to vasculature and blood vessel development (**Fig.S3A**).

We then explored whether there was overlap of the top pathways identified above. Indeed, our analyses indicated significant overlap between immune-related and vascular pathways (**Fig.S3B**). Using Ingenuity Pathway Analysis (IPA), we determined the top upstream regulators predicted to be activated or inhibited based on their respective z-scores. Consistent with enrichment of pro-inflammatory pathways, the top activated upstream regulators in DLPFC were TNF, IL1B, and NFκB complex (**Fig.S3C**). Surprisingly, we discovered that all of the predicted upstream regulators that were inhibited have key roles in the regulation of the vasculature, including KLF2 (163), NRAS (164), and PTEN (165) (**Fig.S3C**). These findings suggest that vascular integrity may be compromised in DLPFC of OUD subjects, in line with growing evidence that the blood brain barrier (BBB) is eroded as consequence of repeated opioid use (166, 167).

As a complementary approach to IPA, we used Hypergeometric Optimization of Motif Enrichment (HOMER) (140) to predict transcription factors that may cause the differential expression of transcripts in OUD. In DLPFC, the top transcription factors were SRF, MEF2A, PKNOX1, PBX3, and SOX21 (**Fig.S3C**). Each of these have roles in neuronal survival (168-176), neuronal activation (177, 178), synaptic plasticity (169, 170), as well as in inflammation (29, 179, 180).

In the NAc, we found the top enriched pathways were related to ECM, including those associated with chondroitin/dermatan sulfate metabolism (**Fig.S4A**). Intriguingly, our data suggests links between ECM remodeling, microglial cell migration, and synaptic plasticity, in line with recent work (181). Specifically, we discovered enrichment of pathways in synapse organization (**Fig.S4A**). We also found enrichment of ameboidal-type cell migration pathways (**Fig.S4A**), commonly used by activated microglia in response to neuroinflammation (182). Whereas there was convergence of cooperative clusters of transcripts into a few, select final common pathways in the DLPFC (**Fig.S3B**), pathways in the NAc were largely discrete (**Fig.S4B**). Using IPA, we predicted the top upstream regulators either activated or inhibited in NAc. The top activated upstream regulators were IL1B (activation of microglia inflammatory response (183)), BRD7 (synaptic plasticity (184)), CREBBP (activity-dependent synaptic remodeling and plasticity (185)), and OSM (cell-type dependent regulation of myelination and inflammation (186)) (**Fig.S4C**). While IKZF1 was predicted to be activated in NAc, this regulator was predicted to be inhibited in DLPFC of OUD subjects, suggesting opioids differentially impact IKZF1 activity and target transcripts in a region-specific manner. Interestingly, IKZF1 is a master regulator of immune cell activation (187-189). The top inhibited upstream regulators were UHRF2 (modulator of DNA methylation (190)), IgG (Complex) (immunomodulation (191)), PTEN (vasculature integrity (165, 192)), ZBTB16 (chromatin remodeling via histone deacetylation (193, 194) including in response to drugs of abuse (195)), and RB1 (cell survival and migration (196)).

As in DLPFC, we identified the top transcription factors using HOMER analysis in NAc in OUD. We found several POU family transcription factors (197) (POU2F3, POU2F2, and BRN1), which have critical roles in inflammation (198-200) and neurodevelopment (201-203). Moreover, we identified the nuclear orphan receptor, RORα, which is also important in neuroinflammation (204-206) (**Fig.S4C**). While the DLPFC and NAc vary in how each coordinate their respective signaling pathways, nevertheless, our data demonstrate that both regions share common neuroinflammation and ECM pathways, suggesting the importance of both in alterations related to chronic opioids.

**Figure S1. Principal component analysis (PCA) indicates lack of batch effect for RNA-sequencing runs. A.** PCA plot for DLPFC. **B.** PCA plot for NAc.

**Figure S2. Differentially expressed transcripts between DLPFC and NAc in control subjects. A.** Table reporting number of DE genes between the DLPFC and NAc of control subjects at various significance cutoffs. A permutation test was used to return corrected p-values and Benjamini-Hochberg q-value are reported**. B.** Log_2_FC plotted relative to -log_10_p-value by volcano plot for DE genes in DLFPC compared to NAc. Horizontal dashed lines represent p-value significance cutoff of corrected p<0.01, while vertical dashed lines represent log_2_FC cutoffs of ≤-0.26 or ≥0.26 (FC≥1.2). Red circles represent DE genes that reach significance, log_2_FC, and FDR<0.05 cutoffs. **C.** Heatmap of the top 250 DE genes (corrected p<0.01 and log_2_FC of ≤-0.26 or ≥0.26) clustered by gene. Each column represents a subject (controls, purple; OUD, blue).

**Figure S3. Principal component analysis (PCA) of unaffected comparison and OUD subjects. A.** PCA plot for DLPFC in unaffected comparison subjects. **B.** PCA plot for DLPFC in OUD subjects. **C.** PCA plot for NAc in unaffected comparison subjects. **D.** PCA plot for NAc in OUD subjects.

**Figure S4**. **Top pathways enriched from differentially expressed transcripts in the DLPFC of OUD subjects.** **A.** Top pathways significantly enriched in the DLPFC ranked by -log_10_ p-value. Pathways mainly include pathways related to inflammation and immune function. **B.** Significantly enriched terms based on pathways included in multiple annotated sets (*e.g.,* GO, KEGG, hallmark, etc.) using hypergeometric p-values and enrichment factors. Note inflammatory response and immune function represent majority of pathways, along with several non-overlapping pathways involved in blood vessel development and vascular processes. **C.** IPA predicted upstream regulators known to be involved in immune function. Analysis focused on regulators predicted to be significantly activated or inhibited (z-scores, yellow to blue gradient represents activated or inhibited, respectively). Top predicted transcription factors and promotor binding sequences by HOMER from DE genes in DLPFC. Several of these transcription factors interact with TNF-dependent signaling, including SRF, MEF2A, and PBX3.

**Fig. S5. Top pathways enriched from differentially expressed genes in the NAc of OUD subjects.** **A.** Top pathways significantly enriched in the NAc ranked by -log_10_ p-value. Pathways mainly include pathways related to chondroitin and dermatan sulfate metabolism and synapse organization. **B.** In the NAc, pathways were distinct, with few overlapping nodes and biological categories. **C.** IPA predicted upstream regulators known to be involved in immune function (activated: IL1B, OSM; inhibited: IgG complex), synaptic plasticity and myelination (inhibited: PTEN), autophagy (inhibited: ZBTB16), and epigenetic regulation of transcription, including chromatin remodeling (activated: BRD7, IKZF1, CREBBP; inhibited: RB1) and DNA methylation (inhibited: UHRF2). Analysis focused on regulators predicted to be significantly activated or inhibited (z-scores, yellow to blue gradient represents activated or inhibited, respectively).

**Fig. S6. Overlapping genes between DLPFC and NAc from rank-rank hypergeometic ordering analysis.** **A.** High overlap between DLPFC and NAc among downregulated and upregulated genes. **B.** Low overlap between DLPFC and NAc among genes with opposing patterns of expression (*i.e.,* downregulated in the DLPFC and upregulated in the NAc, and upregulated in the DLPFC and downregulated in the NAc).

**Fig. S7. Top pathways enriched from differentially expressed genes in the DLPFC and NAc of OUD subjects.** All pathways significantly enriched in both the DLPFC and NAc ranked by -log_10_ p-value.

**Fig. S8. Enrichment of chromatin states in differentially expressed genes in the DLPFC and NAc of OUD subjects. A.** Heatmap of chromatin states at transcription start sites based on annotations for corresponding brain regions and enrichment of upregulated and downregulated differentially expressed (DE) genes. Hypergeometric t-tests were used to calculate the observed overlaps based on observed and expected values. *p<0.01; **p<0.001; and ***p<0.0001. Transcription start site active (TssA); Transcription start site flanking active (TssAFlnk); Transcription at gene 5’ and 3’ (TxFlnk); Strong transcription (Tx); Weak transcription (TxWk); Genic enhancers (EnhG); Enhancers (Enh); Zinc finger genes and repeats (ZNF/Rpts); Heterochromatin (Het); Bivalent/poised transcription start site (TssBiv); Flanking bivalent transcription start site or enhancer (EnhBiv); Repressed polycomb (ReprPC); Weak repressed polycomb (ReprPCWk); Quiescent/low (Quies). **B.** Weak polycomb repression diminishes histone marks (*e.g.,* H3K27me3), which would otherwise inhibit transcription at the start site(26). Relative to baseline, the enrichment of upregulated DE transcripts in DLPFC from OUD subjects suggests that opioid induce the activation of otherwise repressed areas of the genome to promote transcription. In NAc, we discovered DE transcripts were enriched for genomic regions marked for a quiescent state at baseline. Quiescent states are characterized by the complete absence of histone marks linked to transcriptional inactivity (26). Similar to DLPFC, such findings suggest opioids activate otherwise inactive genomic regions, resulting in transcriptional upregulation in NAc.

**Fig S9. Enrichment of synapse-related pathways in differentially expressed transcripts significantly associated with cell-type specific markers in neurons.**

**Fig. S10. Top pathways enriched from genes comprising OUD-specific blue module in the DLPFC and forestgreen module in the NAc.** All pathways significantly enriched in the NAc ranked by -log_10_ p-value. Warmer colors indicate higher enrichment. Gray indicates the absence of enrichment for that pathway in the blue or forestgreen modules.

**Supplementary Tables**

**Table S1. Subject information.** See Excel file.

**Table S2.** Table reporting number of DE genes between OUD and control subjects for DLPFC and NAc at various significance cutoffs. A permutation test was used to return corrected p-values and Benjamini-Hochberg q-value are reported. Note the large numbers of genes that are significantly changed in the NAc relative to the DLPFC from OUD subjects.

|  | **q < 0.01** | **q < 0.05** | **q < 0.1** | **Corrected**  **p < 0.01** | **Corrected**  **p < 0.05** |
| --- | --- | --- | --- | --- | --- |
| DLPFC^1^ | 1  (1/0) | 271 (108/163) | 500  (203/297) | 567 (228/339) | 977 (413/564) |
| NAc^1^ | 53  (2/51) | 1037 (150/887) | 1572  (285/1287) | 1305 (221/1084) | 2095 (419/1676) |

^1^Using cutoff of log2FC ≤ -0.26 and ≥ 0.26; (upregulated / downregulated)

**Table S3. Differentially expressed genes by specific cell-type.** Significant differentially expressed genes were only for neurons and microglia. FDR<0.05. See Excel file.

**Table S4.** **Long noncoding RNAs in DLPFC blue module and NAc forestgreen module for control and OUD subjects.** See Excel file.

**Supplementary Data Files**

**Data file S1. Differentially expressed genes between DLPFC and NAc in control subjects.** See Excel file.

**Data file S2. Differentially expressed genes in DLPFC.** See Excel file.

**Data file S3. Differentially expressed genes in NAc.** See Excel file.
