## Supplementary figures and images for "Transcriptional alterations in opioid use disorder reveal an interplay between neuroinflammation and synaptic remodeling"

### Supplemental Figure 1

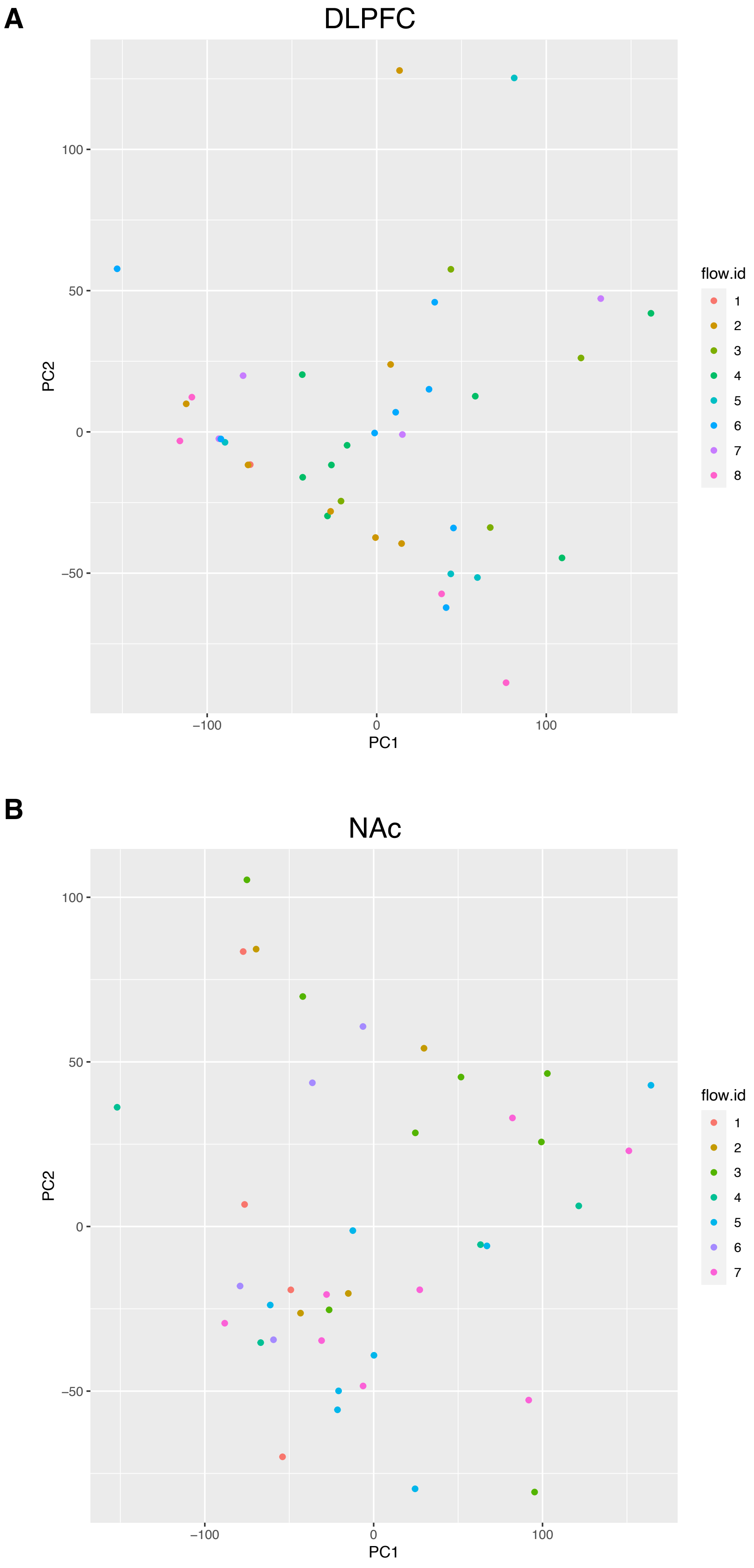

### Supplemental Figure 2

**a**

| | q < 0.01 | q < 0.05 | q < 0.1 | Corrected p < 0.01 | Corrected p < 0.05 |
|---|---|---|---|---|---|
| **DE**[1] | 0 (0/0) | 0 (0/0) | 0 (0/0) | 97 (41/56) | 509 (173/336) |

[1]Using cutoff of $\log_2 FC \leq -0.26$ and $\geq 0.26$; (higher DLPFC / higher NAc)

**b**

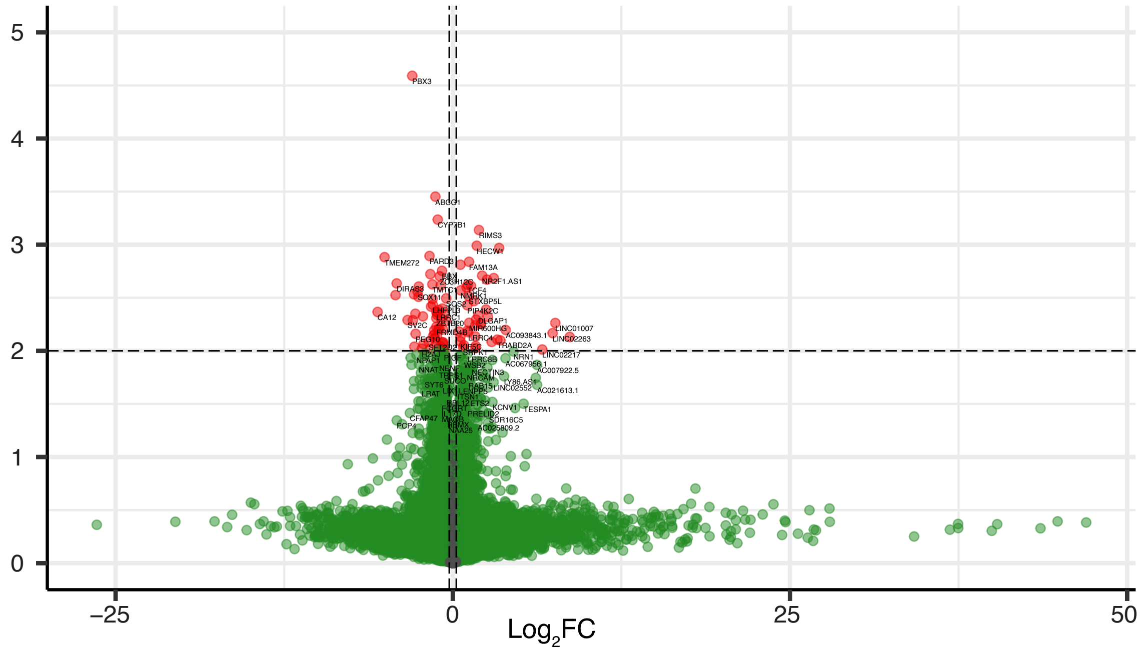


**c**

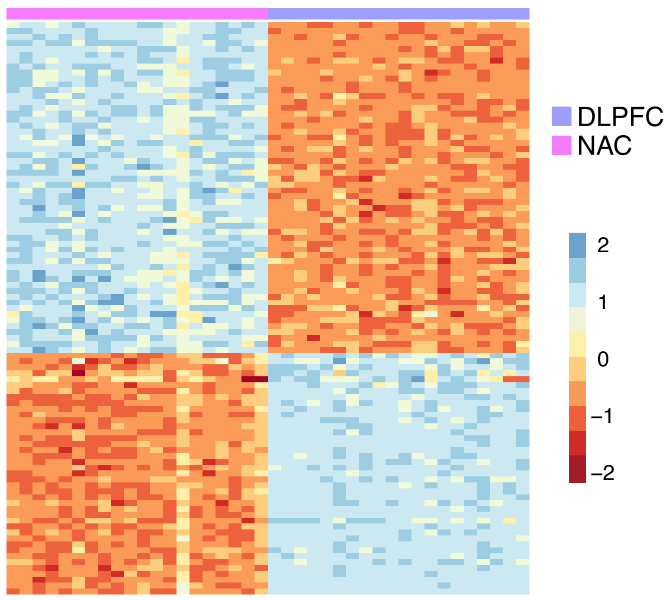

### Supplemental Figure 3

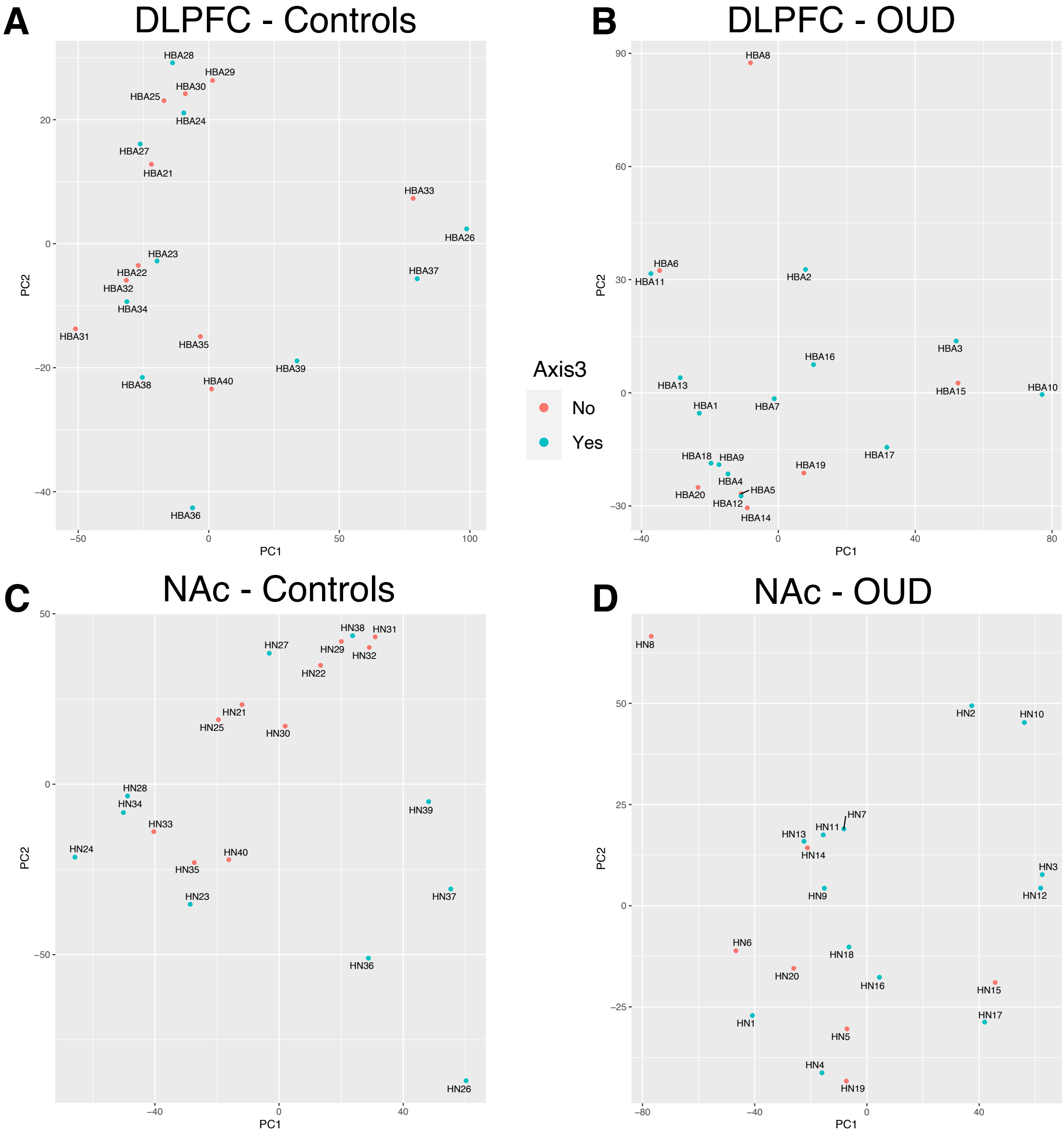

### Supplemental Figure 4

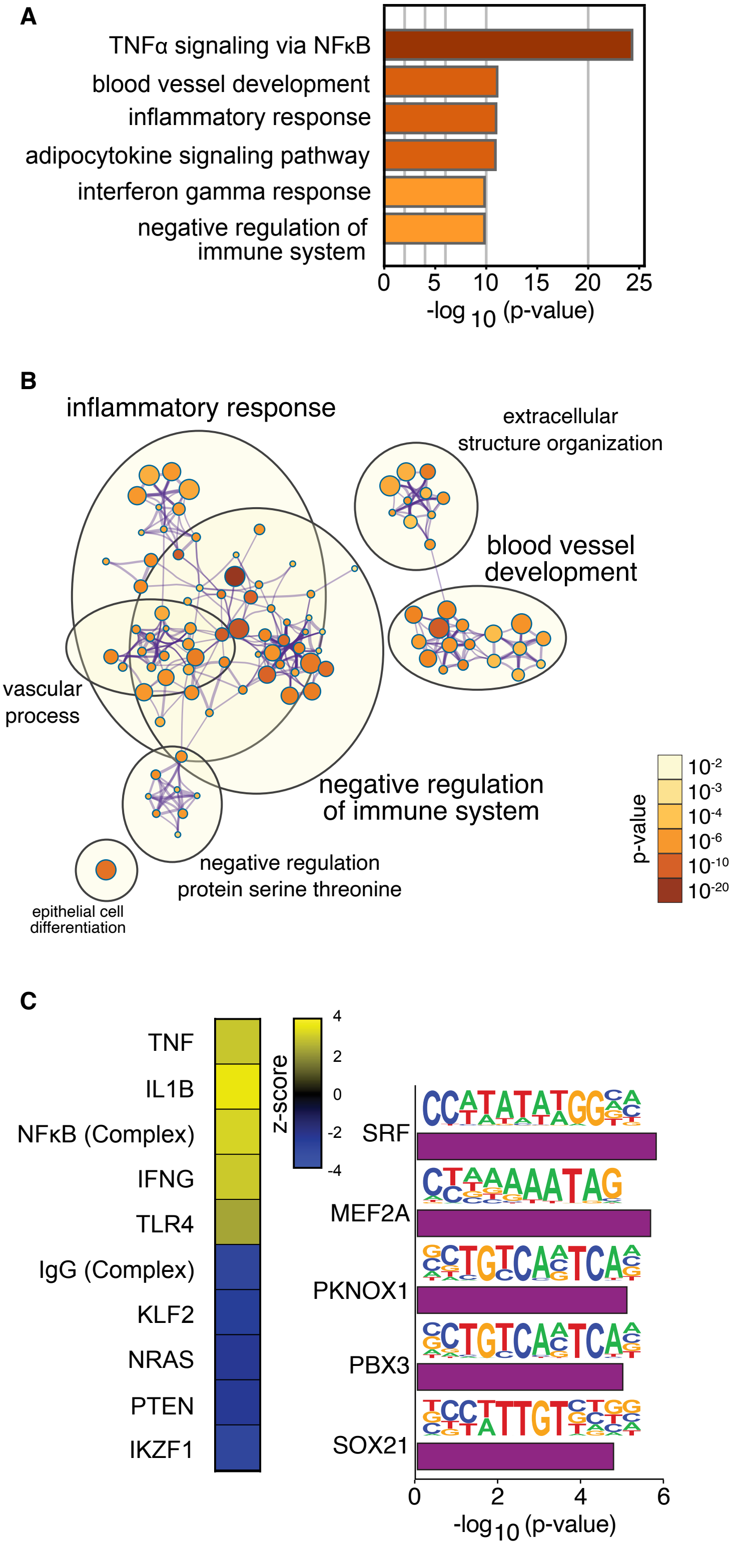

### Supplemental Figure 5

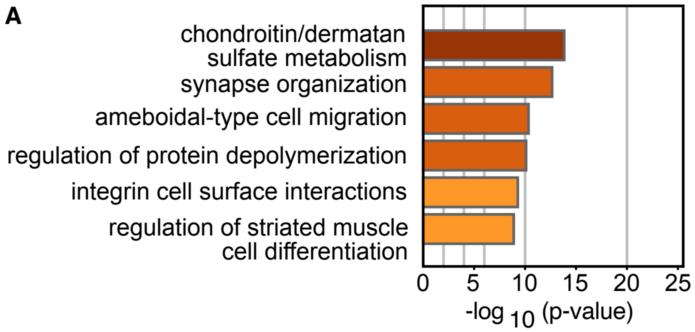

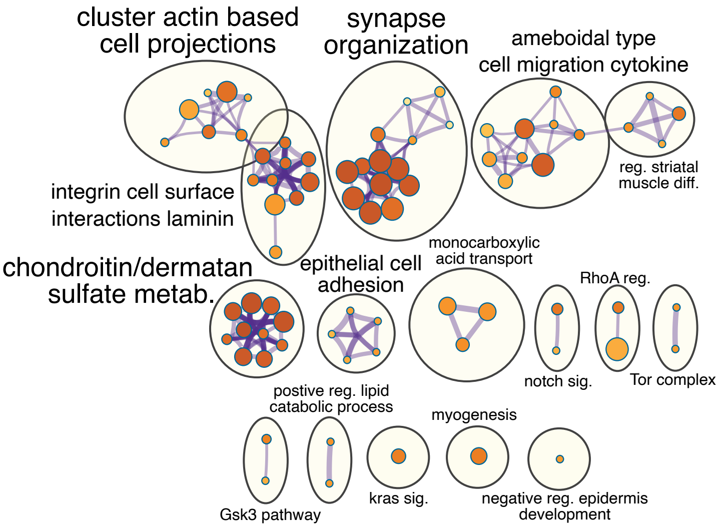

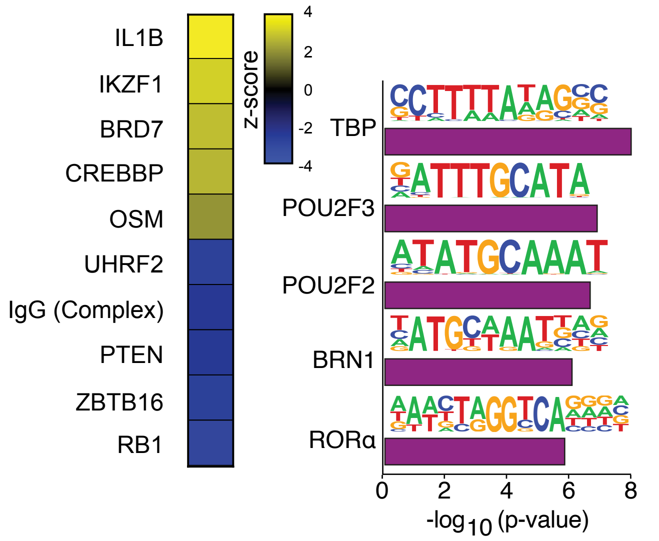

### Supplemental Figure 6

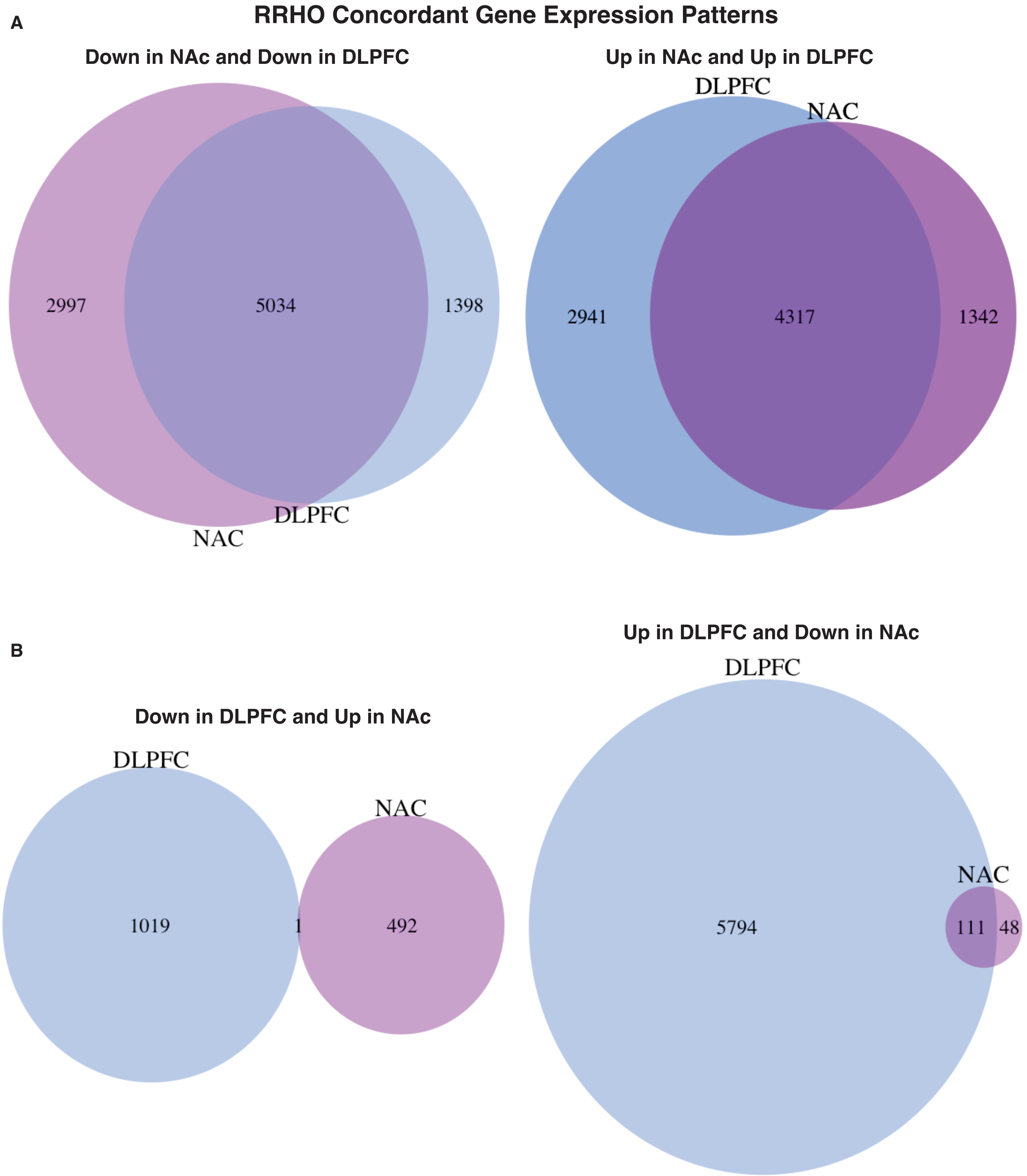

### Supplemental Figure 7

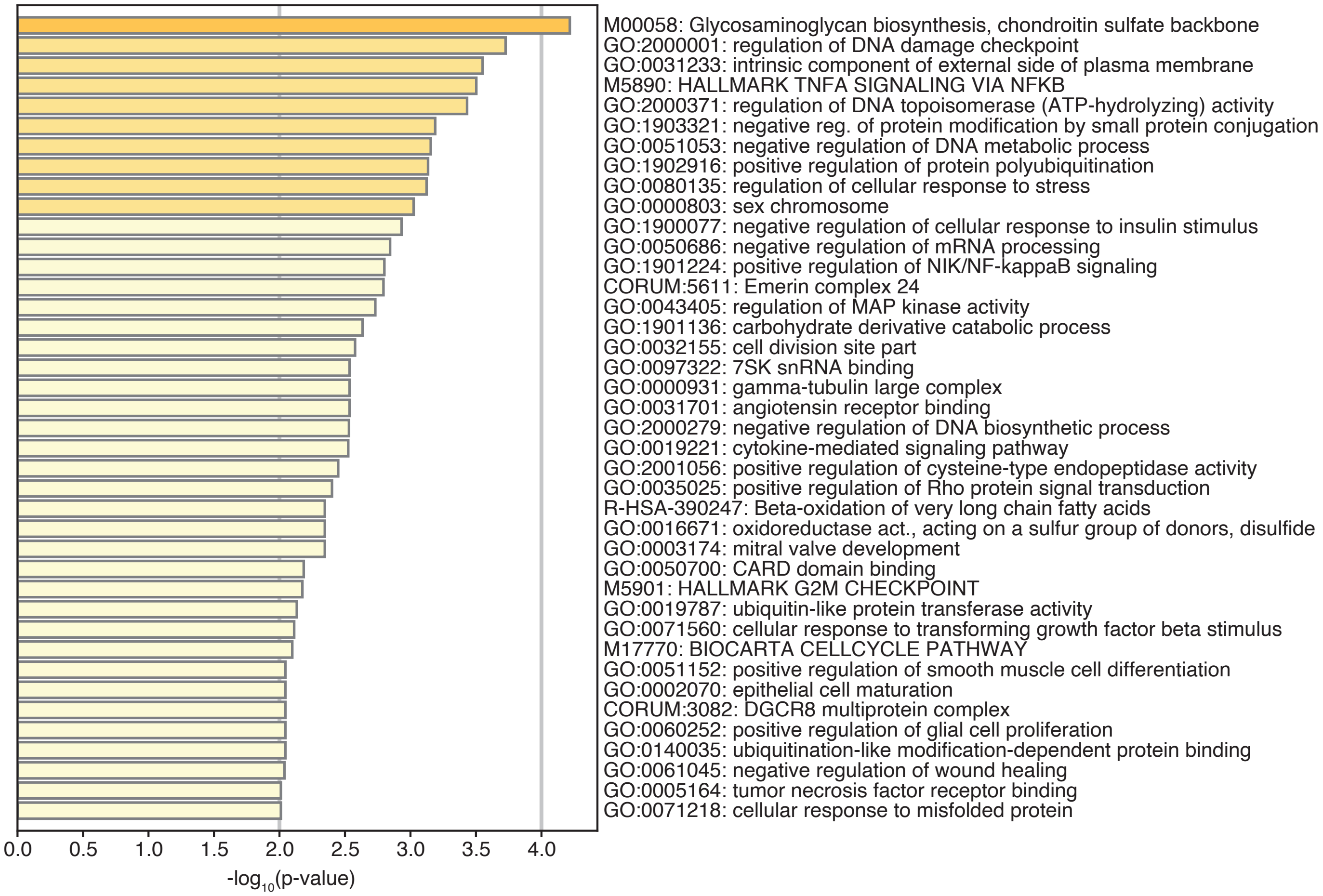

### Supplemental Figure 8

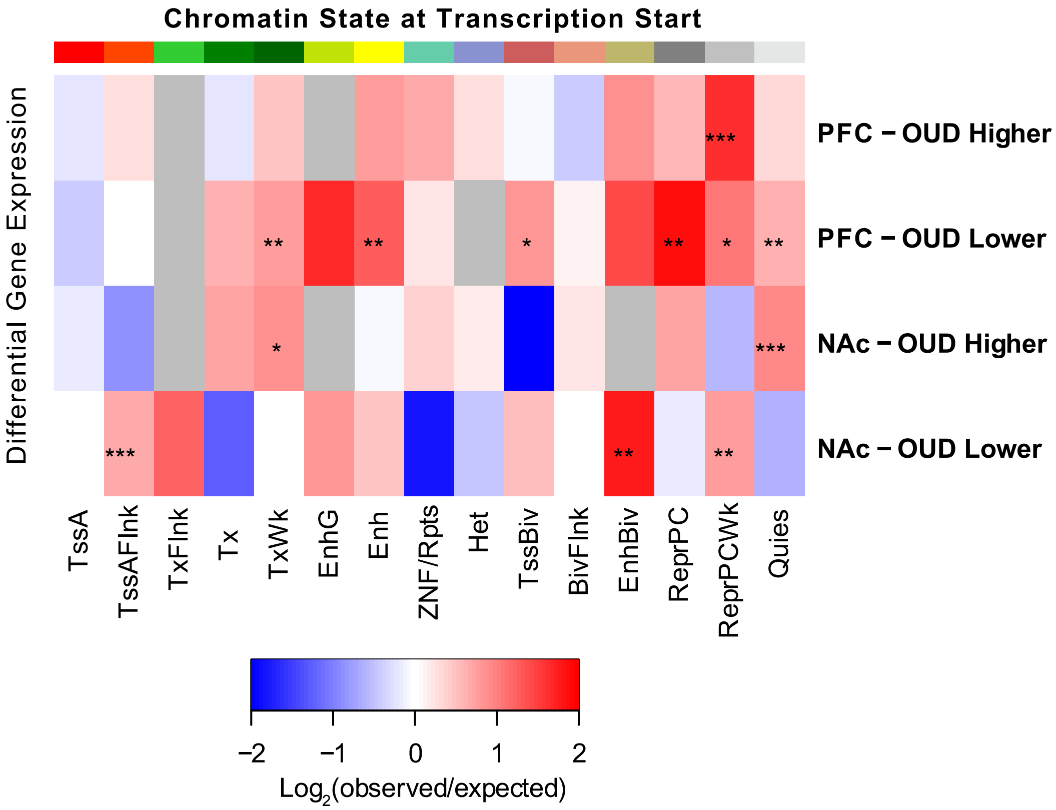

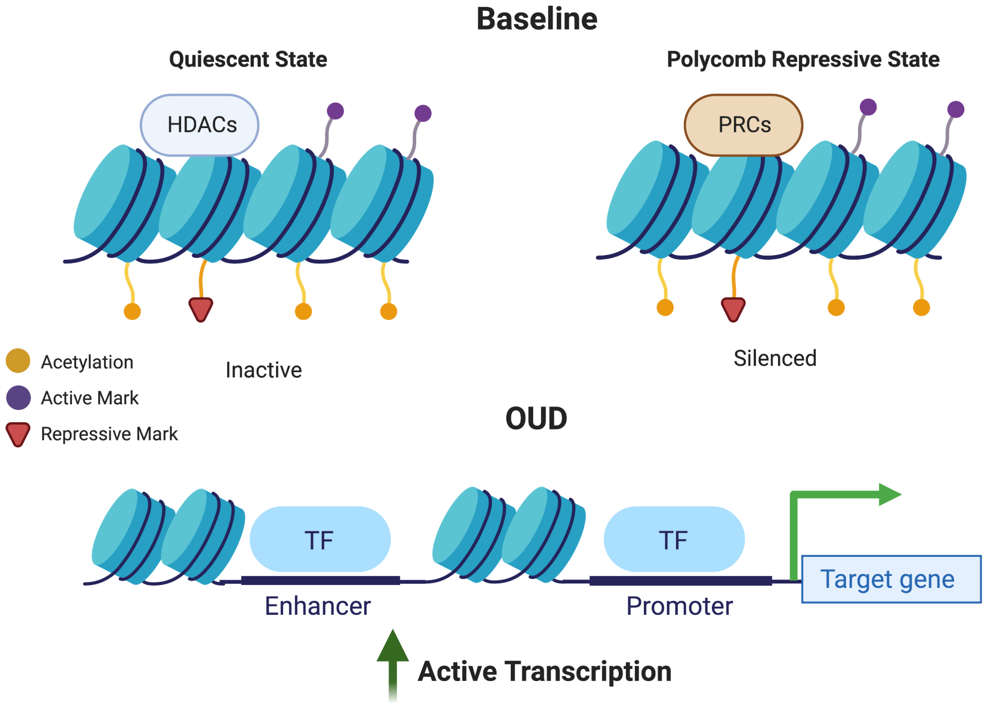

### Supplemental Figure 9

#### regulation of presynapse - synapse organization and assembly

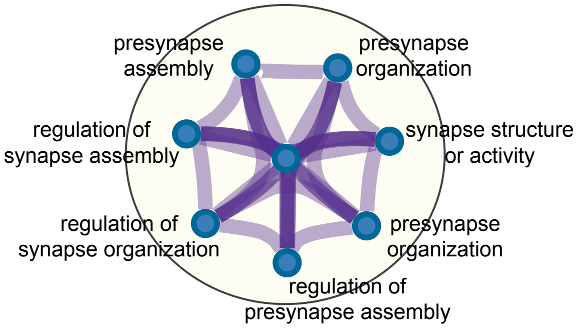

### Supplemental Figure 10

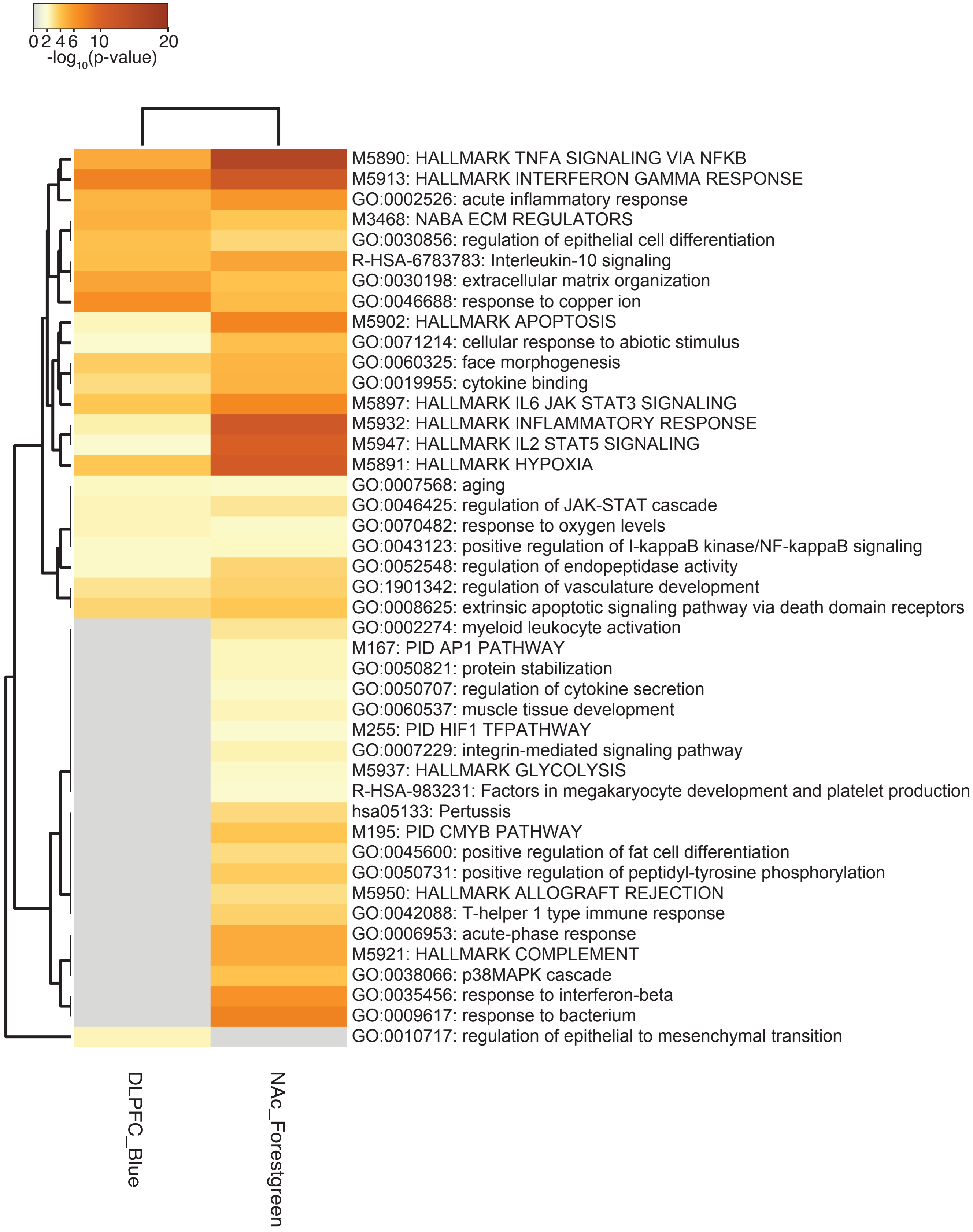
